# Protrudin functions as a scaffold in the endoplasmic reticulum to support axon regeneration in the adult central nervous system

**DOI:** 10.1101/2020.01.15.907550

**Authors:** Veselina Petrova, Craig S. Pearson, James R. Tribble, Andrea G. Solano, Evan Reid, Pete A. Williams, Herbert M. Geller, Richard Eva, James W. Fawcett

**Affiliations:** John van Geest Centre for Brain Repair, Department of Clinical Neurosciences, University of Cambridge, UK; Laboratory of Developmental Neurobiology, Division of Intramural Research, National Heart, Lung and Blood Institute, NIH, Bethesda, USA; Department of Clinical Neuroscience, Division of Eye and Vision, St. Erik Eye Hospital, Karolinska Institutet, Stockholm, Sweden; Cambridge Institute for Medical Research and Department of Medical Genetics, University of Cambridge, UK; Centre for Reconstructive Neuroscience, Institute of Experimental Medicine CAS, Prague, Czech Republic

**Keywords:** Axon regeneration, Protrudin, endoplasmic reticulum, axon transport, optic nerve, neuroprotection

## Abstract

Adult mammalian central nervous system axons have intrinsically poor regenerative capacity, so axonal injury has permanent consequences. One approach to enhancing regeneration is to increase the axonal supply of growth molecules and organelles. We achieved this by expressing the adaptor molecule Protrudin which is normally found at low levels in non-regenerative neurons. Elevated Protrudin expression enabled robust central nervous system regeneration both *in vitro* in primary cortical neurons and *in vivo* in the injured adult optic nerve. Protrudin overexpression facilitated the accumulation of endoplasmic reticulum, integrins and Rab11 endosomes in the distal axon, whilst removing Protrudin’s endoplasmic reticulum localization, kinesin-binding or phosphoinositide-binding properties abrogated the regenerative effects. These results demonstrate that Protrudin promotes regeneration by functioning as a scaffold to link axonal organelles, motors and membranes, establishing important roles for these cellular components in mediating regeneration in the adult central nervous system.

## Introduction

Axons of immature central nervous system (CNS) and adult peripheral nervous system (PNS) neurons readily regenerate after injury (Nicholls and Saunders, 1996; Huebner and Strittmatter, 2009). In contrast, adult CNS neurons lose their regenerative ability with maturation (Bradke and Marín, 2014), meaning that axonal injury or disease has life-altering consequences and that there is little chance of recovery. In addition to the non-permissive extracellular environment after injury, intrinsic neuronal factors also play an important role in the regenerative failure observed in mature CNS neurons (He and Jin, 2016; Tedeschi and Bradke, 2017). Studies aimed at enhancing CNS regeneration have identified transcriptional and epigenetic programs (Gaub *et al*., 2010; Puttagunta *et al*., 2014), signaling pathways (Park *et al*., 2008; Liu *et al*., 2010; Sun *et al*., 2011), the cytoskeleton (Erturk *et al*., 2007; Ruschel *et al*., 2015; Blanquie and Bradke, 2018; Tedeschi *et al*., 2019) and axon transport (Andrews *et al*., 2009; Cheah *et al*., 2016; Eva *et al*., 2017; Koseki *et al*., 2017; Nieuwenhuis *et al*., 2018) as important factors governing regenerative ability. However, the precise machinery required to reconstitute and extend an injured axon is not completely understood, and repairing the injured CNS remains a challenging objective (Fawcett, 2020).

This study focuses on the adaptor molecule Protrudin as a tool for investigating and enhancing axon growth and regeneration in the adult CNS. Protrudin is an integral endoplasmic reticulum (ER) membrane protein that has two properties which make it a candidate for enabling axon regeneration. First, overexpression of Protrudin causes protrusion formation in non-neuronal cell lines, and promotes neurite outgrowth in neuronal cells (Shirane and Nakayama, 2006); second, Protrudin is a scaffolding molecule which possesses interaction sites for key axon growth-related molecules and structures (Shirane and Nakayama, 2006). Through its Rab11 and kinesin-1 (KIF5) binding sites, Protrudin can enable the anterograde transport of Rab11-positive recycling endosomes, leading to their and their cargo’s accumulation at protrusion tips (Shirane and Nakayama, 2006; Raiborg *et al*., 2015). This is relevant to CNS axon repair because increased Rab11 transport into CNS axons *in vitro* increases their regenerative ability (Koseki *et al*., 2017).

Protrudin localizes to the ER through two transmembrane domains and a hairpin loop and interacts with VAP proteins at ER-membrane contact sites through an FFAT domain. This interaction is involved in its effects on protrusion outgrowth (Saita *et al*., 2009; Gil *et al*., 2012; Chang, Lee and Blackstone, 2013). In addition to its localization at ER tubules, Protrudin also regulates ER distribution and network formation (Chang, Lee and Blackstone, 2013). Protrudin also has a FYVE domain that binds to phosphoinositides enabling interaction with endosomes and the surface membrane (Gil *et al*., 2012). Protrudin therefore links a number of cellular components associated with axonal growth. Our hypothesis was that expression of an active form of Protrudin would enable regeneration of CNS axons via a combination of these interactions and would be a powerful tool for understanding the mechanisms of axon regeneration.

Our initial studies found that Protrudin mRNA is expressed at low levels in CNS neurons, but at higher levels in regenerating PNS neurons, and the protein is present in immature regenerative CNS axons but is depleted from axons with maturity as regeneration is lost. We reasoned that overexpression might allow for increased availability of regenerative machinery within the axon, leading to better regenerative after injury. Because stimulation of protrusions through the interaction of Protrudin and Rab11 is increased by growth factor receptor phosphorylation, and its protrusive effects are prevented by dominant negative mutations of the ERK phosphorylation sites, we created a phosphomimetic, active form of Protrudin, mutating these previously identified phosphorylation sites (Shirane and Nakayama, 2006).

Here, we report that Protrudin promotes axon regeneration through the mobilization of endosomes and ER into the distal part of injured axons, whilst having striking effects on neuronal survival after axotomy. Importantly, Protrudin expression only has moderate effects on developmental axon growth but has strong effects on regeneration. Overexpression of Protrudin, particularly in its active form, led to robust regeneration of mature cortical axons *in vitro* and of retinal ganglion cell (RGC) axons in the injured optic nerve. Protrudin expression caused increased transport of Rab11 endosomes, integrins, and an accumulation of ER in the axon tip, with active Protrudin increasing this effect. Deleting either the ER transmembrane domains or VAP-binding FFAT domain prevented the accumulation of the ER whilst also abrogating the effects on regeneration. Interfering with other key domains of also eliminated the regenerative effects, indicating that Protrudin promotes regeneration by acting as a scaffold in the axonal ER, bringing together growth components, organelles and membranes to enable CNS axon regeneration.

## Results

Protrudin has several key domains, including a Rab11-binding domain (RBD), three hydrophobic membrane-association domains (TM 1-3), an FFAT motif for binding to VAP proteins at ER contact sites, a coiled-coiled (CC) domain (which interacts with kinesin 1) and a phosphoinositide-binding FYVE domain which enables interaction with endosomes and the plasma membrane (Shirane and Nakayama, 2006; Saita *et al*., 2009; Matsuzaki *et al*., 2011; Gil *et al*., 2012; Chang, Lee and Blackstone, 2013). The map of these domains is shown in **Fig. 1A**. **Fig. 1B** shows the phosphorylation sites that were mutated to produce active Protrudin in accordance with previous literature (Shirane and Nakayama, 2006).

**Fig. 1.**
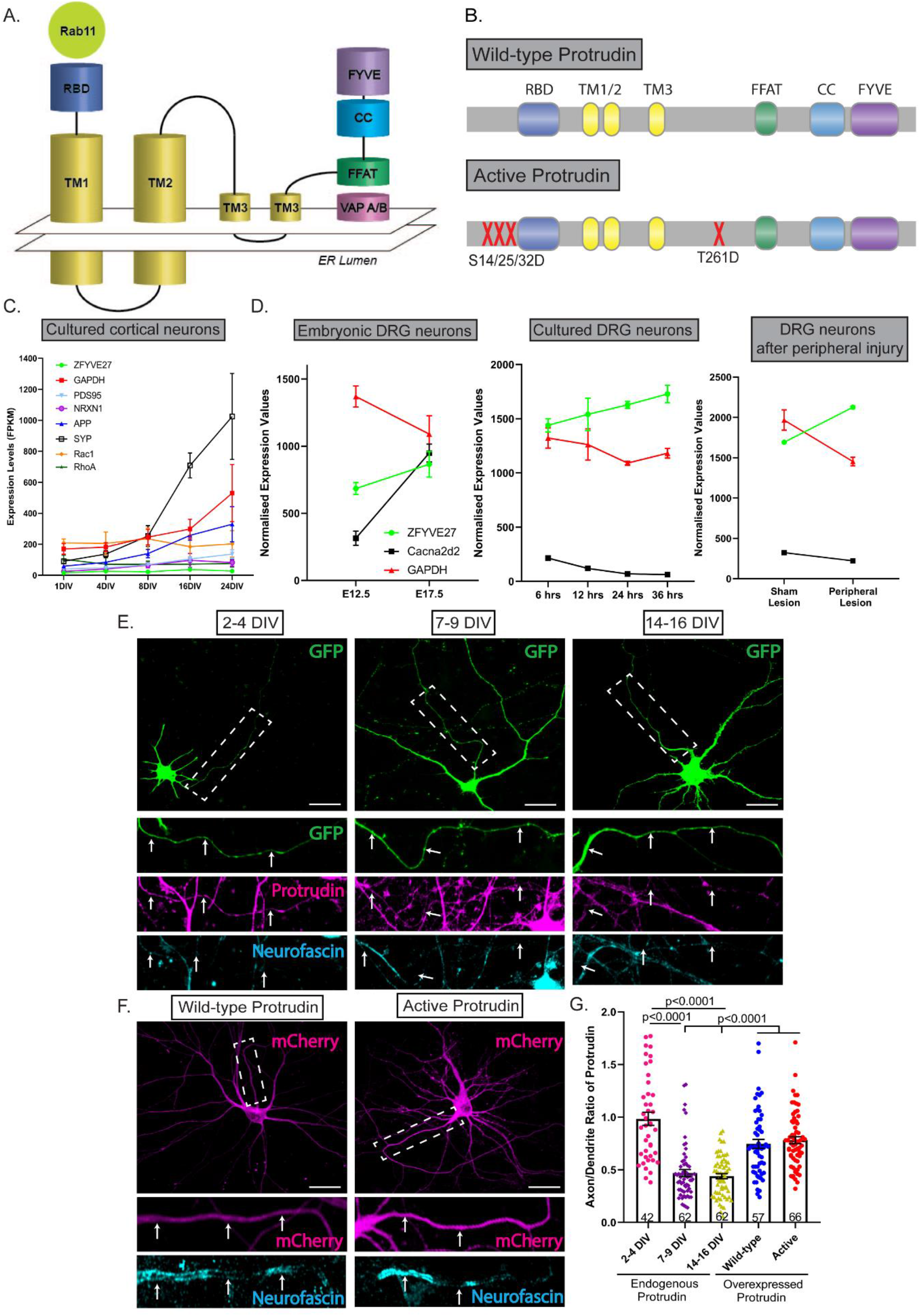
Protrudin is expressed at low levels in mature axons and overexpression restores this deficit. (**A**) Schematic diagram of Protrudin’s domains and structure. (**B**) Schematic of wild-type and active Protrudin mutagenesis sites. (**C**) mRNA expression levels of six neuronal genes (including *Zfyve27 –* the Protrudin gene) from different stages of development in primary rat cortical neurons *in vitro*. (**D**) Normalized expression levels of Protrudin and other related genes during embryonic development in the mouse, after plating DRG neurons *in vitro* or after peripheral nerve injury in DRG cells. (**E**) Immunofluorescent images of Protrudin in the proximal axons (white dotted line box) of neurons at different stages of development in culture. Scale bars are 20 µm. The white arrows follow the course of the proximal axon. (**F**) Immunofluorescent images of overexpressed, mCherry-tagged wild-type or active Protrudin (magenta) and staining for the axon initial segment marker – neurofascin (cyan). Scale bars are 20 µm. (**D**) The axon-to-dendrite ratio of Protrudin at different developmental stages or after overexpression (*n* = 4, *Kruskal-Wallis with Dunn’s* multiple comparison test, *p* < 0.0001). Error bars represent mean ± SEM.

### Protrudin is expressed at low levels in mature, non-regenerative CNS neurons

In order to affect growth and regeneration, we reasoned that Protrudin would need to be present in axons in significant quantity and be optimally functional. We first examined the mRNA expression of Protrudin in developing CNS and PNS neurons, as well in PNS neurons after injury, using previously published RNA-seq datasets. We found that Protrudin mRNA (*Zfyve27*) is expressed at low levels in CNS neurons, and its expression is not developmentally regulated (**Fig. 1C**) (Koseki *et al*., 2017). In contrast, in sensory, regeneration-capable neurons, the Protrudin transcript increases with development, during axon growth *in vitro*, and in response to peripheral nerve injury (**Fig. 1D**) (Tedeschi *et al*., 2016).

To assess the level and distribution of Protrudin protein in CNS axons, we examined its endogenous localization in rat primary cortical neurons by immunocytochemistry. We compared developing neurons (2-4 days after plating at E18), with mature neurons that have lost the ability to regenerate their axons (matured *in vitro* for 14+ days). We found that Protrudin localized to both axons and dendrites of developing cortical neurons but was enriched in dendrites and restricted from axons at later stages, coinciding with the time when cortical neurons mature and lose their regenerative ability (**Fig. 1E**) (Koseki *et al*., 2017). By measuring the axon and dendrite fluorescence intensity of Protrudin immunolabelling we found that younger neurons (2-4 days *in vitro*, DIV) had a higher axon-to-dendrite ratio (ratio=1) compared with later stages of development (7-9 DIV, ratio=0.47 and 14-16 DIV, ratio=0.44) (**Fig. 1G**). These observations suggest that endogenous Protrudin may not be present in sufficient quantity in mature CNS axons to influence regeneration. Overexpression of either wild-type or active Protrudin resulted in a substantial increase in the protein level in rat primary cortical neurons (**Fig. S1**). This resulted in an increased axon-to dendrite ratio in neurons overexpressing wild-type (ratio=0.77) or active Protrudin (ratio=0.78) indicating an increase in the protein’s distribution to axons, with Protrudin easily detectable throughout axons (**Fig. 1F-G**). The exclusion of Protrudin from mature axons is therefore not absolute and can be overcome by overexpression. Overexpression of Protrudin had no effect on soma size or spine number and morphology whilst active Protrudin had a modest effect on increasing dendritic tree complexity (**Fig. S2**).

### Overexpression of wild-type or active Protrudin enhances axon regeneration *in vitro*

Protrudin overexpression has previously been associated with enhanced neurite outgrowth in HeLa and PC12 cells as well as primary hippocampal neurons at early stages of development (Shirane and Nakayama, 2006). Given that Protrudin is expressed at low levels in CNS neurons, we reasoned that its overexpression might increase axon growth. We transfected wild-type or active Protrudin into immature 2 DIV cortical neurons and measured the growth of axons and dendrites at 4 DIV (**Fig. 2A**). Overexpression of active Protrudin had a modest effect on this early phase of axon growth, with neurons overexpressing active Protrudin having increased length of the longest neurite (580 µm) compared to control-transfected neurons (470 µm) (**Fig. 2B**). Overexpression of wild-type Protrudin had no effect. These results show that high levels of active Protrudin have a small effect on the initial outgrowth phase of neurons in culture. At this time axon growth is already rapid, axons usually regenerate when cut, and the maturation-related compartmentalization of the neurons to exclude growth-related molecules from axons has not yet occurred (Koseki *et al*., 2017).

**Fig. 2.**
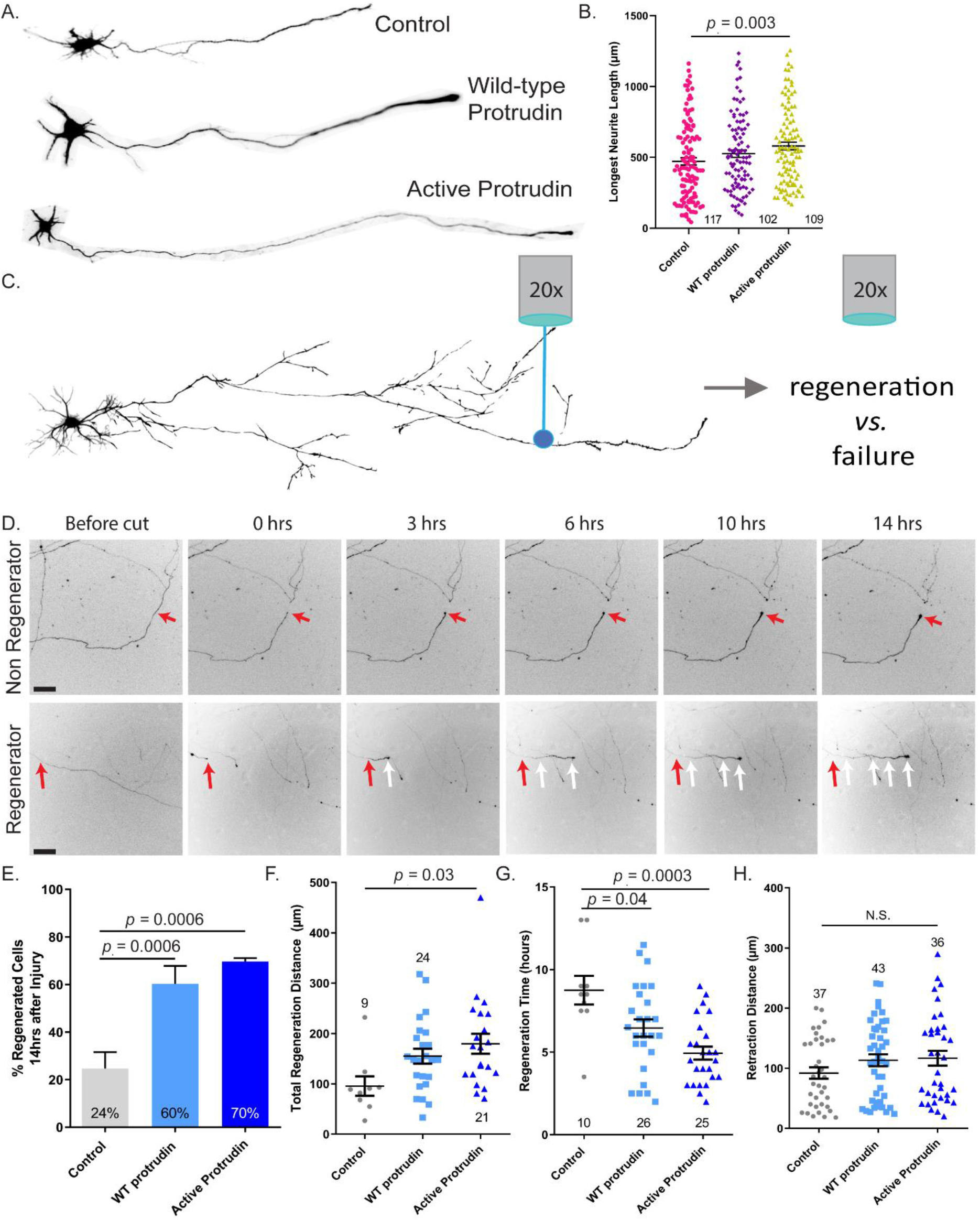
Protrudin overexpression has a modest effect on initial neurite outgrowth but is a strong promoter of axon regeneration after laser axotomy. (**A**) Example neurons at 4 DIV overexpressing control construct, wild-type or active Protrudin. (**B**) The average length of the longest neurite in each condition (*n* = 3, *p* = 0.01, *Kruskal-Wallis with Dunn’s* multiple comparison test). Error bars represent mean ± SEM. (**C**) Diagram of the laser axotomy method (**D**) Representative images show a regenerating and a non-regenerating axon over 14 h post laser axotomy. The red arrows at 0 h post injury shows the point of injury. The white arrows trace the path of a regenerating axon. Scale bars are 50 µm. (**E**) Percentage of regenerating axons overexpressing either mCherry control (*n* = 45), mCherry wild-type Protrudin (*n* = 45) or mCherry active Protrudin (*n* = 39) (*Fisher’s exact* test). (**F**) Quantification of regeneration distance 14 h after injury (*One-way ANOVA*, *p* = 0.04). (**G**) Quantification of regeneration initiation time (*One-way ANOVA*, *p* = 0.0004). (**H**) Quantification of retraction distance (*One-way ANOVA*, *p* = 0.214). Error bars represent mean ± SEM.

We next examined the effect of Protrudin overexpression on axon regeneration in mature axons. We used the laser axotomy model of regenerative decline, where axons progressively lose their regenerative ability as they mature and become electrically active (Eva *et al*., 2017; Koseki *et al*., 2017)(**Fig. 2C**). Cortical neurons were again transfected with either control, wild-type or phosphomimetic, active Protrudin, this time at 10 DIV. We examined axon regeneration after laser injury at 13-17 DIV, when regenerative capacity has declined (Koseki *et al*., 2017). In this model, axons typically show two responses to injury, either the formation of a retraction bulb and no regeneration, or retraction and bulb formation followed by growth cone development and axon extension (**Fig. 2D**). Expression of either wild-type or active Protrudin led to a dramatic increase in the percentage of axons regenerating after laser axotomy (**Fig. 2E, Video S1**) with axons regenerating longer distances (**Fig. 2F**) and initiating regeneration in a shorter time (**Fig. 2G**). These regenerative events were most pronounced in neurons transfected with active Protrudin. No differences in the retraction distance after injury were observed (**Fig. 2H**). Importantly, overexpressed wild-type and active Protrudin was found to localize throughout axons, accumulating at the growth cones of uninjured axons, at regenerating growth cones, and at the retraction bulbs of non-regenerating injured axons. At the growth cone, Protrudin localized principally to the central domain (**Fig. S3D-F**).

Protrudin’s effect on axon regeneration was dose dependent; co-transfection with a construct encoding GFP resulted in lower Protrudin expression and a reduced effect on regeneration (**Fig. S3A-C**). These results show that Protrudin, particularly in its constitutively active phosphomimetic form, has a very strong effect on the rescue of axon regeneration in mature neurons. There is therefore a contrast between Protrudin’s minor enhancement of the outgrowth of immature axons that are already growing rapidly, and the rescue of regeneration in mature neurons whose axons seldom regenerate.

### Protrudin promotes regeneration through increased endosomal transport

Protrudin has several interaction sites that have the potential to link endosomes, ER, membrane and kinesin. Our hypothesis, based on studies of how Protrudin causes neurite growth in HeLa and PC12 cells (Shirane and Nakayama, 2006) and our studies of Rab11 vesicles and their cargo in regeneration (Koseki *et al*., 2017), was that the main effect of Protrudin would be to enable transport of Rab11 vesicles and their contents into mature axons through linkage to KIF5, so increasing regenerative capacity. In order to determine if Protrudin’s regenerative effects were mediated through enhanced axonal transport, we examined the transport of Rab11 in the presence of wild-type or active Protrudin, and also studied the transport of a known Rab11 cargo, integrin alpha 9 (Eva *et al*., 2010). This integrin can mediate long range sensory regeneration in the spinal cord (Cheah *et al*., 2016). In addition, we overexpressed three mutated Protrudin constructs targeting domains associated with endosomal transport (Rab-binding domain, KIF5-interaction domain, and FYVE domain), and examined axon regeneration after laser axotomy.

To determine whether Protrudin’s regenerative effects were also accompanied by an increase in axonal transport, we used spinning-disc live-cell microscopy to observe the movement of Rab11-GFP or integrin α9-GFP in the distal part of mature, 13-17 DIV axons in the presence of overexpressed wild-type or active Protrudin (**Fig. 3A**). Vesicle transport was scored as anterograde, retrograde, bidirectional or static and the total number of Rab11 or integrin α9-positive endosomes per section of axon was measured. The majority of Rab11-positive vesicles trafficked bidirectionally whereas the bulk of integrin-containing endosomes moved retrogradely confirming previous studies (Eva *et al*., 2010, 2017). Overexpression of wild-type or active Protrudin resulted in increased retrograde and bidirectional transport of Rab11-GFP and enhanced anterograde and retrograde transport of integrin α9-GFP (**Fig. 3B-D**), leading to more total Rab11 and integrin-positive vesicles in the distal axon (**Fig. 3B-D**). Approximately 20% of α9-transporting vesicles were positive for Protrudin (**Fig. S4A-B**) and 29% of Protrudin-positive endosomes (wild-type or active) were also Rab11-positive (**Fig. S4A-B**), and kymograph analysis demonstrated dynamic co-localization of both Rab11 and α9-integrin with Protrudin.

**Fig. 3.**
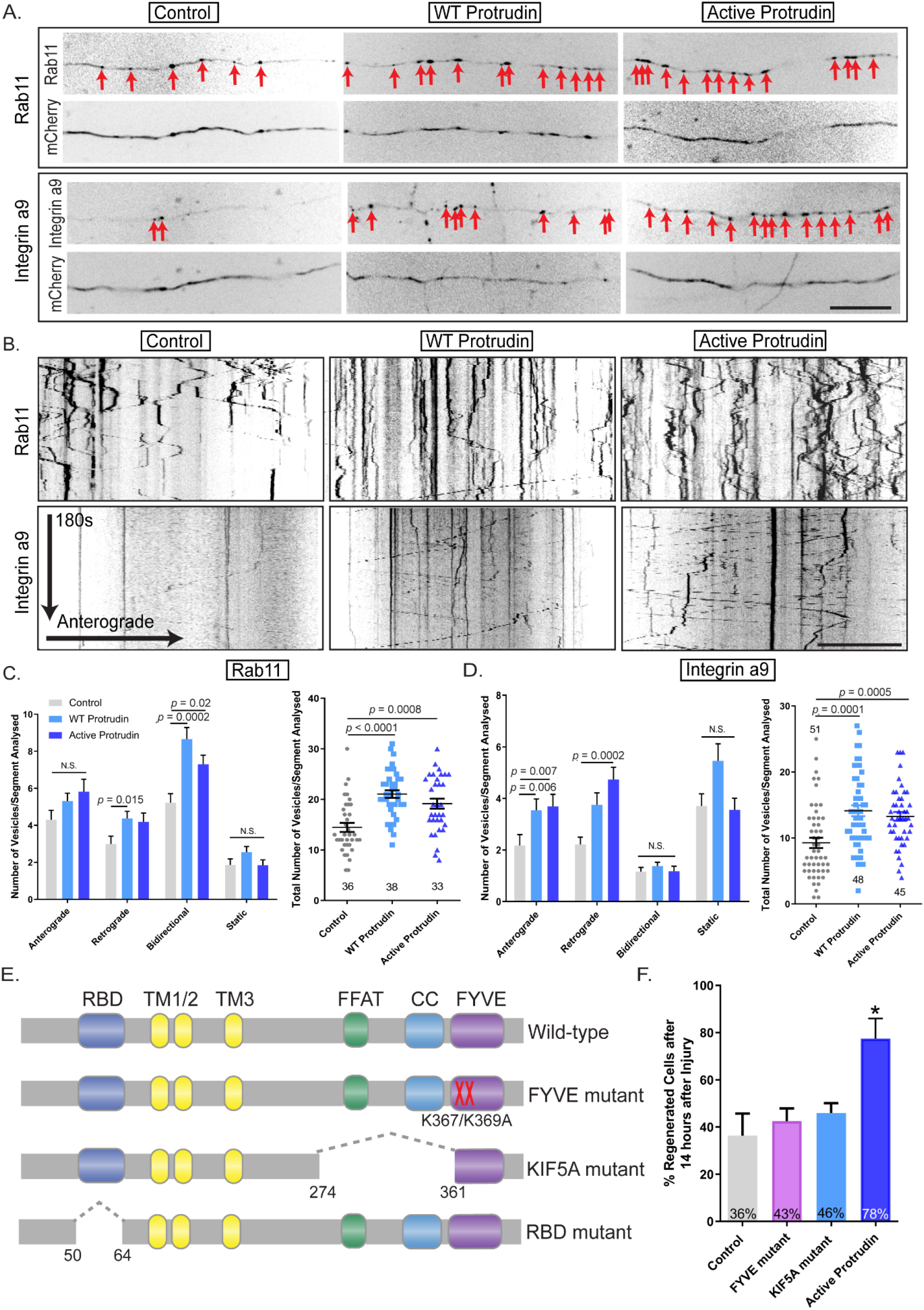
Protrudin enhances the transport of growth machinery and receptors in the distal axon, and its involvement in axon transport is required for axon regeneration. (**A**) Representative distal axon sections of neurons expressing integrin α9-GFP or Rab11-GFP, together with either mCherry (control), mCherry-wild-type Protrudin or mCherry-active Protrudin. (**B**) Kymographs showing the dynamics of integrin α9-GFP and Rab11-GFP in distal axons of co-transfected neurons. Scale bar is 10 µm. (**C**) Quantification of Rab11-GFP axon vesicle dynamics and total number of Rab11 GFP vesicles in distal axon sections (*Kruskal-Wallis with Dunn’s* multiple comparison test) (**D**) Quantification of integrin α9-GFP axon vesicle dynamics and total number of integrin α9-GFP vesicles in distal axon sections (*Kruskal-Wallis with Dunn’s* multiple comparison test). Error bars represent mean ± SEM. (**E**) Schematic representation of Protrudin transport domain mutants. (**F**) Percentage of regenerating axons in neurons expressing mCherry-Protrudin domain mutants – FYVE (*n* = 56) and KIF5A (*n* = 56) compared to active Protrudin as a positive control (*n* = 24), and mCherry as a negative control (*n* = 42).

To test the contribution of Protrudin’s transport-associated domains towards its regenerative effects, we assembled a cohort of mutants in accordance with previous literature (Shirane and Nakayama, 2006; Matsuzaki *et al*., 2011; Gil *et al*., 2012). We deleted the Rab-binding domain (RBD) which is required for Rab11 anterograde transport (Shirane and Nakayama, 2006). We also mutated the coiled-coil domain which is essential for the interaction with the anterograde axonal motor KIF5 (Matsuzaki *et al*., 2011) and made a dominant negative FYVE domain mutant which prevents the interaction of Protrudin with phosphoinositides on endosomal membranes (Gil *et al*., 2012) (**Fig. 3E**). Each of these mutants was separately expressed in cortical neurons at 10 DIV and their effects on the regeneration of mature axons were quantified at 13-17 DIV using laser axotomy. Unexpectedly, overexpression of the RBD mutant caused extensive neuronal cell death (**Fig. S4C-D**), indicating an essential role for the Protrudin-Rab11 interaction in neuronal viability but precluding examination of its effect on axon regeneration. Mutations of the KIF5 or the FYVE domain sharply diminished the effects of Protrudin overexpression on axon regeneration, whilst active Protrudin again stimulated robust regeneration (**Fig. 4F**). These data demonstrate that the interactions between Protrudin, endosomes and KIF5 are required for the axon regeneration-promoting effects of Protrudin, and that Protrudin-driven regeneration is accompanied by an increase in the anterograde transport of integrins and Rab11 endosomes.

**Fig. 4.**
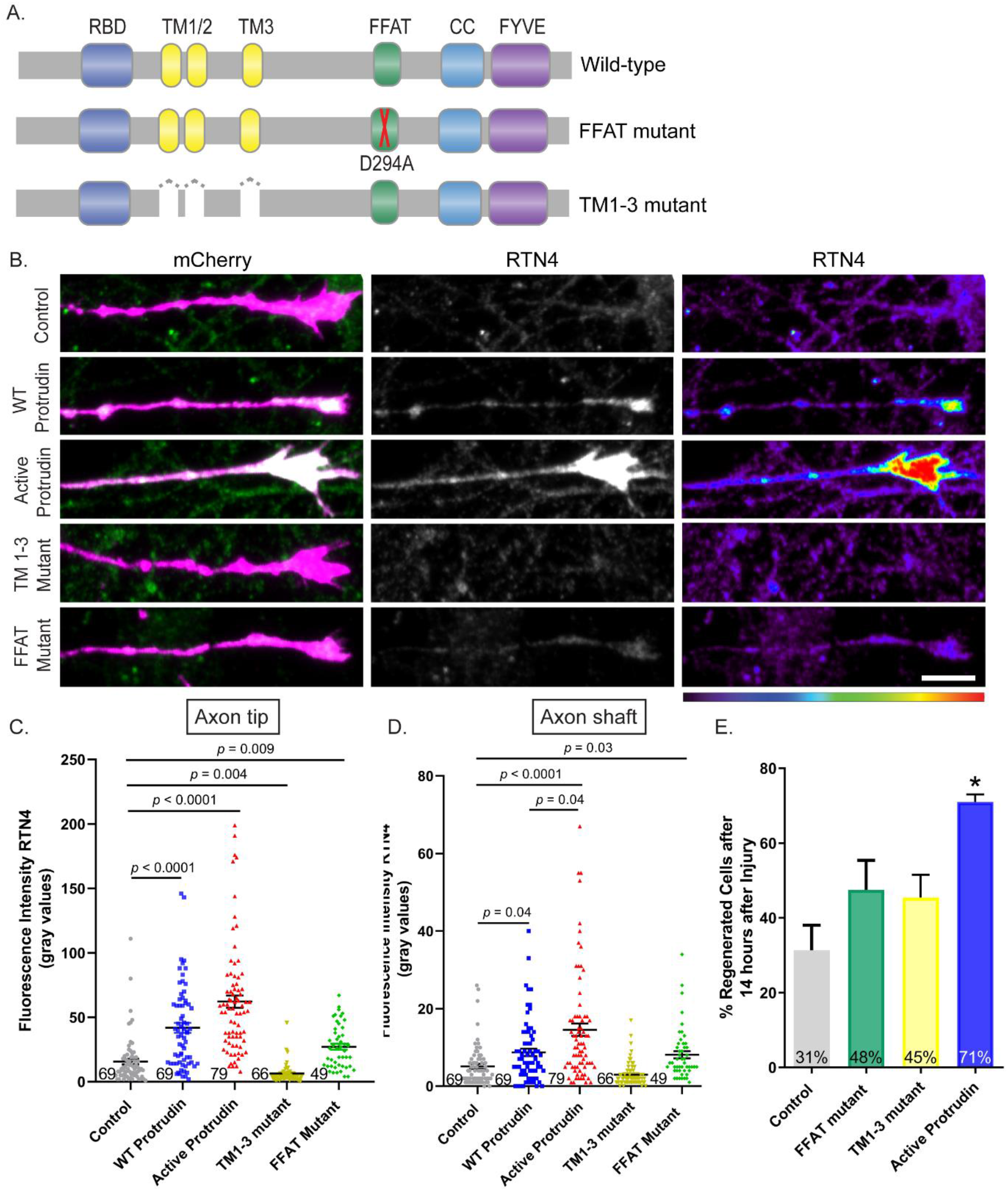
Protrudin overexpression enhances ER presence at growth cones and this interaction is required for successful axon regeneration. (**A**) Schematic representation of Protrudin endoplasmic reticulum domain mutants. (**B**) Representative images of RTN4 immunofluorescence (green) in the distal axon of neurons expressing the indicated m-Cherry Protrudin constructs (magenta). Scale bar is 10 μm. (**C-D)** Quantification of RTN4 fluorescence intensity at the axon tip and shaft (*p* < 0.0001, *Kruskal-Wallis with Dunn’s* multiple comparisons test). Error bars represent mean ± SEM. (**E**) Percentage of regenerating axons in neurons expressing mCherry-Protrudin domain mutants – FFAT (*n* = 60) and TM1-3 (*n* = 45) compared to active Protrudin as a positive control (*n* = 21), and mCherry as a negative control (*n* = 41).

### Protrudin promotes regeneration through interaction with the endoplasmic reticulum

There is increasing evidence that endosomal transport is heavily influenced by ER-endosome contact sites, and that the distribution and morphology of ER tubules is controlled by kinesin-dependent endosomal transport (Raiborg *et al*., 2015; Lee and Blackstone, 2020). Additionally, ER-endosome and ER-plasma membrane contact sites have been observed in both axons and dendrites (Wu *et al*., 2017). We reasoned that the localization of Protrudin to the ER, and its interaction with contact site proteins might contribute to its regenerative effects. In order to study this, we mutated the FFAT domain which is important for Protrudin’s interaction with VAP proteins at ER contact sites (Chang, Lee and Blackstone, 2013), and we created a mutant lacking all three transmembrane (TM1-3) domains which confer its membrane localization within the ER (**Fig. 4A**). Deletion of these hydrophobic regions releases Protrudin from the ER, rendering it cytosolic (Chang, Lee and Blackstone, 2013).

The ER exists as a continuous tubular organelle through axons (similar to an axon within the axon), and its genetic disruption causes axonal degeneration (Yalçın *et al*., 2017). Re-establishment of the axonal ER may be equally as important as the re-establishment of the axon membrane for successful regeneration. Because ER tubules undergo highly dynamic movements, partly by hitchhiking on motile endosomes (Guo *et al*., 2018), we hypothesized that linkage of overexpressed Protrudin to kinesin might lead to an increase in tubular ER in the axon. To examine the effects of Protrudin overexpression on axonal ER we analyzed the distribution of reticulon 4, which reports on ER abundance in axons (Farías *et al*., 2019) (**Fig. 4B**). Overexpression of wild-type and active Protrudin resulted in increased reticulon 4 in the growth cone shaft and at the axon tip of uninjured axons (**Fig. 4B-D**). Crucially, mutation of FFAT or TM1-3 domains abolished this effect.

In order to study the importance of the Protrudin-ER interaction for Protrudin-mediated axon regeneration, each of the mutants described above was separately expressed in primary rat cortical neurons at 10 DIV and their effects on axon regeneration were studied at 13-17 DIV using laser axotomy. Both mutants sharply diminished the effects of Protrudin overexpression on axon regeneration, whilst active Protrudin again stimulated robust regeneration (**Fig. 4E**). This finding demonstrates that Protrudin can carry ER into axon growth cones, and that this ER localization is necessary for Protrudin’s regenerative effects. This indicates an important role for the ER in mediating CNS axon regeneration.

Collectively, the results so far demonstrate that Protrudin enables axon regeneration by acting as a scaffold that links key players that participate in regeneration. Axonal ER, recycling endosomes, kinesin 1, and phosphoinositides, are all brought together in distal axons and regenerating growth cones. The finding that mutation of any of the binding domains in Protrudin abrogates its effect on regeneration suggests that the co-location of all these components is necessary for efficient axon regeneration.

### Overexpression of wild-type or active Protrudin promotes axon regeneration in the injured optic nerve

Protrudin’s robust effect on CNS axon regeneration *in vitro* prompted us to investigate its effectiveness on optic nerve regeneration. We first examined Protrudin mRNA levels in retinal ganglion cells (RGCs) in published RNA-seq datasets (Williams *et al*., 2017) and found that Protrudin mRNA is present at low levels in mature, adult RGCs (**Fig. 5A**). This corresponded with our findings in cortical neurons but not in regenerative PNS neurons where Protrudin levels are much higher (**Fig. 1C-D**). We generated three constructs for AAV delivery to the retina by intravitreal injection: AAV2-GFP, AAV2-ProtrudinGFP, and AAV2-activeProtrudin-GFP. The viruses transduced 40-45% of RGCs throughout the retina and the protein was observed throughout uninjured axons (**Fig. 5C-D**). Higher Protrudin levels were detected by immunohistochemistry of wholemount retinas in eyes following overexpression the wild-type or active Protrudin but not after control virus (**Fig. S5A**). Viruses were injected 2 weeks before optic nerve crush, and 2 weeks after injury the anterograde axon tracer – cholera toxin subunit-ß (CTB) was administered (**Fig. 5B**). Control optic nerves had limited regeneration (0% >0.5 mm from the crush site), while regenerating axons extended to 2.75 mm for wild-type Protrudin-transduced animals, and 3.5 mm for active Protrudin-transduced animals. The numbers of regenerating axons were high, particularly for active Protrudin, in which over 630 axons were seen proximally, significantly more than in control (44 axons) or in wild-type Protrudin (380 axons) (**Fig. 5E-F**). Co-localization between CTB and GAP43 was found throughout the nerve in all conditions suggesting that the majority of CTB-positive axons observed in the nerve past the injury site are regenerating axons (**Fig. S5B**). These findings confirm that Protrudin overexpression leads to a substantial increase in axon regeneration in the injured CNS.

**Fig. 5.**
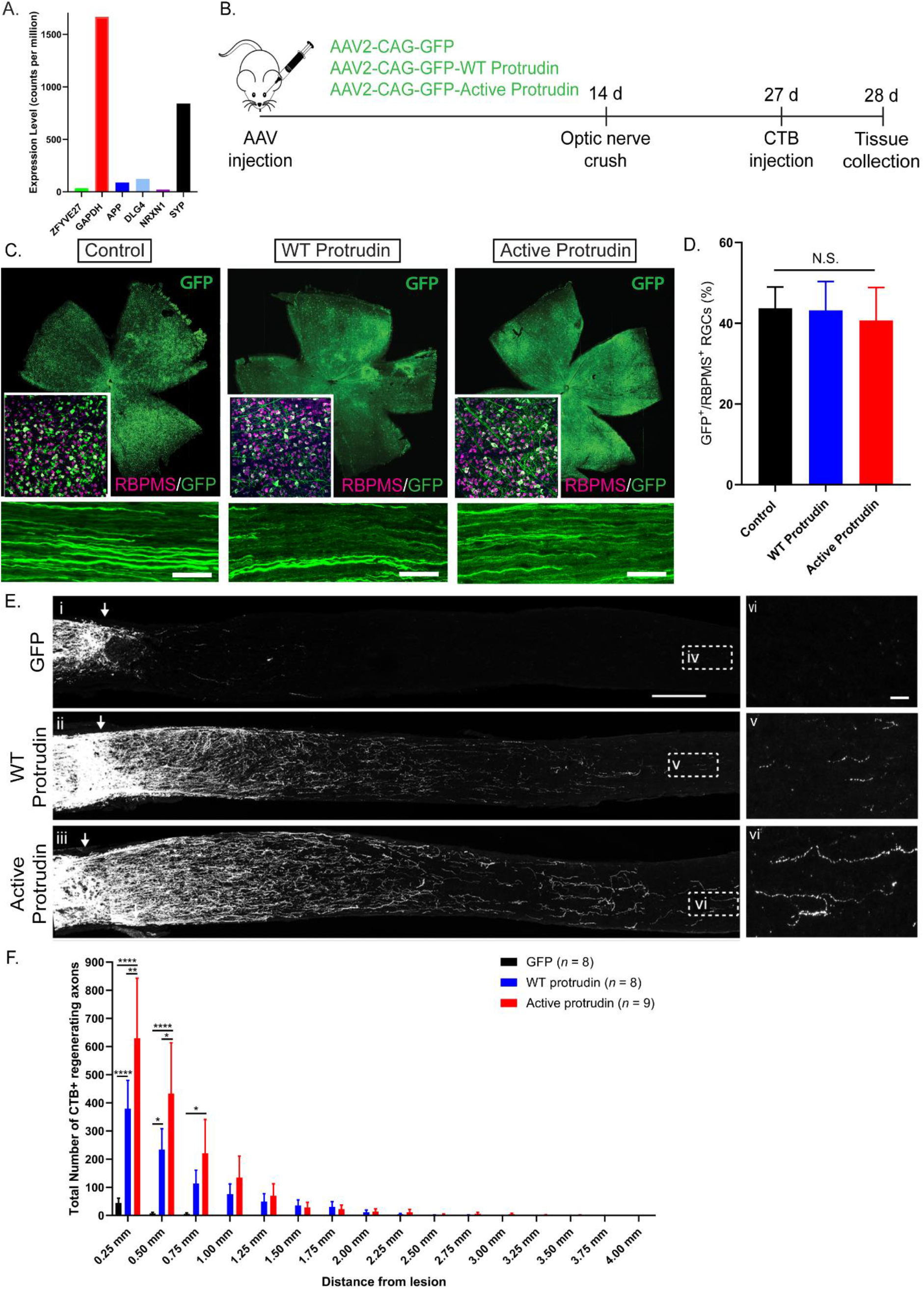
Protrudin enhances regeneration of RGC axons after optic nerve crush. (**A**) Protrudin mRNA levels during the progression of glaucoma in comparison to other neuronal markers. (**B**) Experiment timeline for optic nerve crush. (**C**) Retinal wholemounts or axonal sections showing GFP-positive cells which have been transduced with one of three viruses: control GFP, wild-type Protrudin or active Protrudin-GFP. Retinas were immunostained for RBPMS - retinal ganglion cell marker (magenta) showing co-localization between the virally infected cells (GFP) and RGCs. Scale bar is 20 µm in axonal sections. (**D**) Robust viral expression in RGC-positive cells was detected for all constructs (*n* = 2 per condition). (**E**) CTB-labelled axons in the optic nerves of mice transduced with viruses for wildtype (WT) Protrudin, active Protrudin and GFP control. Arrows indicate lesion site. Insets (iv-vi) show regenerating axons in the distal optic nerve. Scale bar is 200 μm and on inset is 20 μm (*n* = 8-9 animals/group). (**F**) Quantification of regenerating axons at increasing distances distal to the lesion site, displayed as mean ± SEM. Statistical significance was determined by *two-way ANOVA with Bonferroni post-hoc* test for multiple comparisons. ** *p* < 0.005, *** *p* < 0.001, **** *p* < 0.0001

### Overexpression of wild-type or active Protrudin is neuroprotective *in vivo*

We next examined the effects of Protrudin expression on RGC survival. Because optic nerve injury leads to severe neuronal loss two weeks after injury (typically 80-90%) we used an acute retinal explant model which is often used to detect potential neuroprotective treatments for glaucoma (Bull *et al*., 2011; Pattamatta, McPherson and White, 2016). Viruses were injected intravitreally, and retinas were removed two weeks later and cultured as explants for three days (**Fig. 6A**). Both wild-type and active Protrudin were entirely neuroprotective, with these retinas exhibiting no loss of RGC neurons, whilst GFP-only controls lost 55% of their RGCs (**Fig. 6B-C**). Additionally, both Protrudin constructs showed widespread general neuroprotection as there was no reduction in the DAPI-positive cells after injury (**Fig. 6D**).

**Fig. 6.**
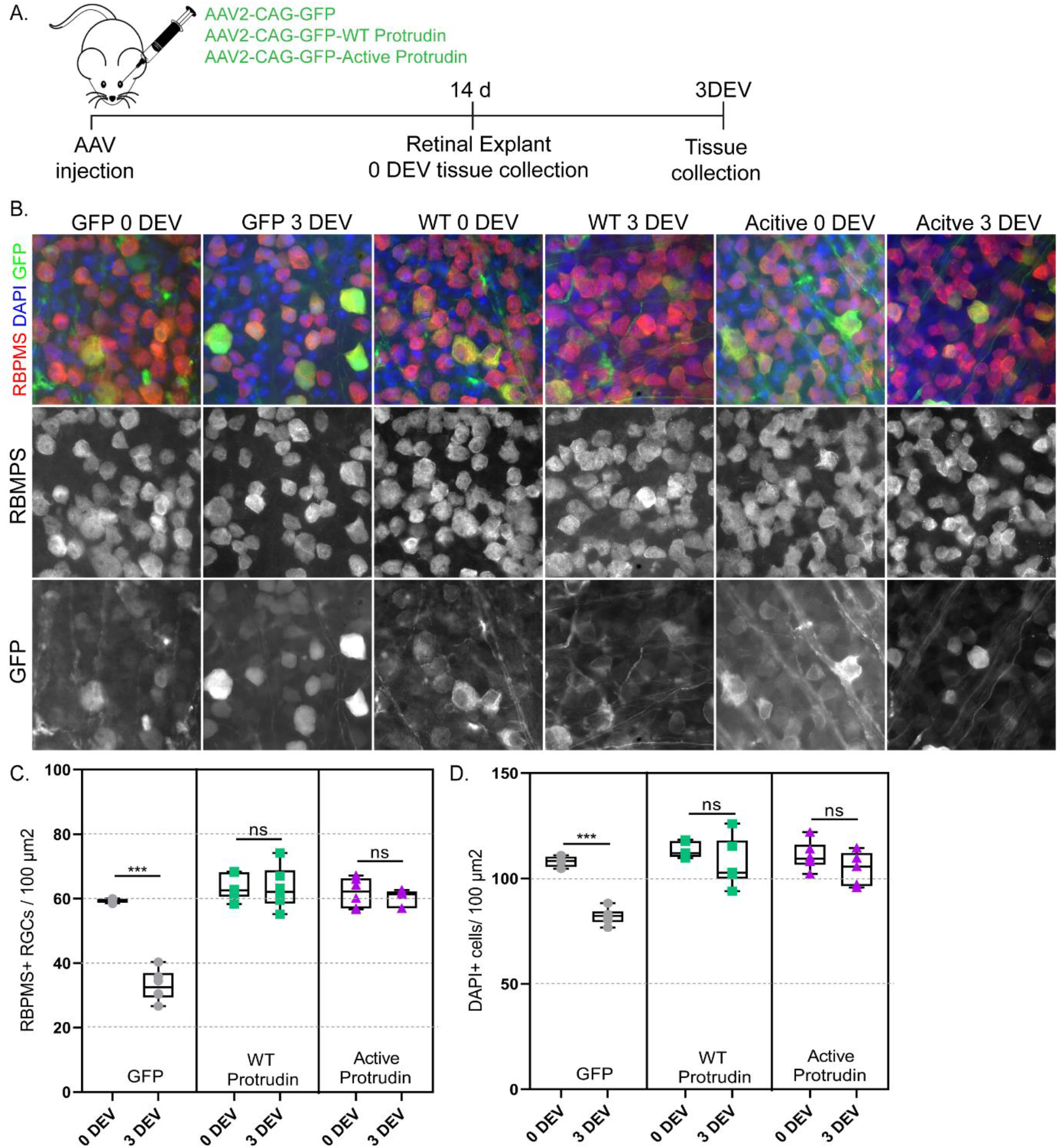
Protrudin is neuroprotective to RGCs and other cell types in the retina after a retinal explant. (**A**) Experimental timeline for retinal explant experiment. (**B**) Representative images of RGCs (red for RBPMS) 0 and 3 days *ex vivo* (DEV) in eyes injected with control virus, wild-type Protrudin or active Protrudin (green for GFP) and stained for DAPI (blue). (**C**) Quantification of RGC survival in retinal explant (*Student’s* t-test). ** *p* < 0.005, *** *p* < 0.001, **** *p* < 0.0001. (**D**) Quantification of DAPI-positive cell survival in retinal explant (*Student’s* t-test). ** *p* < 0.005, *** *p* < 0.001, **** *p* < 0.0001.

## Discussion

Our study demonstrates axon regeneration in the CNS driven by overexpression of the scaffold molecule, Protrudin. Protrudin enables robust axon regeneration and neuroprotection in the retina and optic nerve and promotes regeneration after axotomy of cortical neurons *in vitro.* The action of Protrudin is to bind recycling endosomes, endoplasmic reticulum and kinesin and carry them and their contents to the tip of axons and growth cones.

Protrudin mRNA is expressed at low levels in cultured CNS cortical neurons compared to other abundantly expressed proteins but is found at much higher levels in sensory neurons (**Fig. 1C-D**). Interestingly, Protrudin’s pattern of expression in the developing mouse embryo as well as in cultured DRG neurons and after peripheral lesions contrasts with that of Cacna2d2 – a calcium channel protein which when suppressed improves axon regeneration (**Fig. 1D**) (Tedeschi *et al*., 2016). Protrudin’s upregulation after peripheral nerve lesion suggested pro-regenerative properties.

In previous reports, Protrudin has been detected in mouse primary hippocampal neurons at 1 DIV predominantly localized to the pericentrosomal compartment and to growing neurites, and present in dendrites and at the growth cone (Shirane and Nakayama, 2006). Here, we examined the distribution of the endogenous protrudin protein in rat primary cortical cultures over development from 2-4 DIV to maturation at 14-16 DIV at which time the neurons are spontaneously electrically active and have lost much of their ability to regenerate their axons (Koseki *et al*., 2017). We found that Protrudin’s distribution changes with neuronal maturity – at immature stages (2-4 DIV), the molecule is equally distributed between axons and dendrites, but at later stages (7-9 DIV and 14-16 DIV), axonal levels fall while dendritic levels are maintained (**Fig. 1E-G**). These results suggest that early in CNS neuronal development Protrudin is present in the newly extending axons but then as neurons mature and polarize, the Protrudin protein is redistributed towards the cell body and dendrites and is not available to participate in regeneration. Many molecules become selectively excluded from axons and directed to dendrites as neurons mature and compartmentalize. Previously we have studied the selective distribution of Rab11 and integrins which are essential for neurite outgrowth and axon transport (Andrews *et al*., 2009; Koseki *et al*., 2017). For these molecules the exclusion from axons is more complete than for Protrudin which is driven in axons when the levels in the cell body are raised by overexpression. The exclusion of growth-related molecules from mature axons is one of the reasons for their failure to regenerate, and restoration of Rab11 vesicles and integrins to mature axons can restore regeneration (Andrews *et al*., 2009, 2016; Koseki *et al*., 2017).

The main aims of this study were to determine whether Protrudin can enable axon growth and regeneration and investigate its mechanism of action. Previously, the ability of Protrudin to stimulate process outgrowth was shown to be dependent on its phosphorylation. Because receptors able to activate proteins by phosphorylation may be sparse in mature CNS axons (Petrova and Eva, 2018), a construct for phosphomimetic constitutively active Protrudin was made, based on the previously identified phosphorylation sites (Shirane and Nakayama, 2006).

Previous reports have shown that overexpression of Protrudin can enhance neurite outgrowth in PC12 cells and in hippocampal neurons at early stages of development (Shirane and Nakayama, 2006). We first overexpressed Protrudin during the initial outgrowth period in cultures of immature cortical neurons. Dissociation removes the processes from cells, and after plating they rapidly grow axons and then dendrites. Overexpression of Protrudin during this rapid growth phase led to a modest increase in axon length when active Protrudin when overexpressed in rat cortical neurons (**Fig. 2A-B**). However, we did not observe any differences when wild-type Protrudin was overexpressed as predicted by previous studies in hippocampal neurons (Shirane and Nakayama, 2006).

The next step was to measure the effects of Protrudin on axon regeneration. For this, we used a culture model in which neurons mature over time, and the probability of axons regenerating declines from around 70% to 5% after 20DIV (Koseki *et al*., 2017), reproducing the decline in intrinsic regenerative ability that is seen with neuronal maturity *in vivo.* Our *in vitro* regeneration data showed that wild-type and active Protrudin greatly enhanced axon regeneration after laser injury (**Fig. 2C-H**). The percentage of regenerating axons especially in the active Protrudin condition (70%) was found to be higher than some of the best treatments utilised previously in this model system such as depletion of EFA6 – an ARF6 activator in the axon (59%) (Eva *et al*., 2017) and overexpression of dominant negative Rab11 (38%) (Koseki *et al*., 2017). 70% appears to be the ceiling value for regeneration in this assay. Interestingly, it was observed that overexpression of wild-type Protrudin was capable of enhancing axon regeneration although to a slightly lesser extent than active Protrudin. This effectiveness of wild-type Protrudin could be due to phosphorylation of the overexpressed protein by endogenous kinases therefore producing effects similar to active Protrudin.

Once Protrudin’s regenerative ability was confirmed, we studied potential mechanisms, based on Protrudin’s many interaction domains. Protrudin’s actions affected transport as predicted, because overexpression of wild-type and phosphorylated Protrudin resulted in increased transport of Rab11 endosomes and in enhanced anterograde and retrograde transport of integrins in the distal axon (**Fig. 3A-D**). This study focused on integrin α9 because of its ability to promote long-range axon regeneration in the spinal cord and on Rab11 because these endosomes transport integrins (Andrews *et al*., 2009; Cheah *et al*., 2016); however, the effects of Protrudin on axon growth are most likely both integrin dependent and independent (Eva *et al*., 2010). In addition to integrins, Rab11 endosomes transport many growth-promoting molecules that could influence regeneration including the IGF-1 and TrkB receptors (Romanelli *et al*., 2007; Hollis *et al*., 2009a; Hollis *et al*., 2009b). Rab11 may also promote regeneration through regulation of membrane trafficking events (Campa and Hirsch, 2017). In addition, based on the original hypothesis, the domain’s linking axonal transport to membrane and endosomal trafficking (FYVE domain, KIF5A domain and Rab11-binding domain) were mutated, and each of these changes suppressed the axon regeneration-promoting effect of Protrudin (**Fig.3E-F**).

Axons are enriched with ER tubules, whilst ER sheets are predominantly found in cell bodies. The axonal ER is a network of tubules which are highly dynamic and exist as a continuous network throughout the axon (Öztürk, O’Kane and Pérez-Moreno, 2020). This ER tubule continuity is required for axonal integrity and survival (Yalçın *et al*., 2017) and for interaction with microtubules during development to control axon polarity and growth (Farias, 2019). The involvement of the ER in axon regeneration has not been extensively studied in mammalian CNS neurons before. One previous study has presented correlative evidence implicating not only the ER but also the protrudin-binding protein-spastin in axon but not dendrite regeneration due to their accumulation at the tips of regenerating axons after axonal injury in *Drosophila* (Rao *et al*., 2016) and ER-microtubule interaction is involved in the establishment of neuronal polarity (Farías *et al*., 2019). Here, we found that active Protrudin overexpression increased ER (shown by reticulon 4, a marker of smooth ER) (Farías *et al*., 2019) in axonal growth-cones (**Fig. 4B-D**). In addition, TM1-3 mutant Protrudin and FFAT mutant Protrudin, both important for ER localization, each eliminated its ability to enrich ER at growth cones and to promote axon regeneration after laser axotomy (**Fig. 4**). There are several mechanisms by which enrichment of the ER could facilitate axon regeneration, including bulk transfer of lipids from the ER to the plasma membrane, synthesis and transfer of signaling lipids, calcium signaling and involvement in organelle trafficking. ER-plasma membrane interaction sites are a potential site for some of these interactions (Raiborg *et al*., 2015, 2016). Protrudin-mediated enrichment of ER into the tip of growing processes could function to enable the transfer of lipids, with the FYVE domain promoting interaction with the surface membrane (Gil *et al*., 2012), allowing for rapid expansion of the growth cone plasma membrane - a requirement for successful axon regeneration (Bradke, Fawcett and Spira, 2012). Late endosomes in contact with ER, interacting via Protrudin and FYCO1 have been implicated in process outgrowth in PC12 cells (Raiborg *et al*., 2015; Hong *et al*., 2017), but FYCO1 is absent in the rat neurons used for the present study (Koseki *et al*., 2017). A variant of the mechanism probably contributes to Protrudin-mediated outgrowth in cortical neurons.

The overall result of these studies into the interaction domains of Protrudin is that all of them are involved in axon regeneration, and that inactivation of any of them removes most of the ability of Protrudin to promote regeneration. The conclusion is that Protrudin works by bringing all of its binding partners together in such a way that they can collaborate in enabling axon regeneration.

The next step was to examine Protrudin’s effects *in vivo.* The optic nerve crush model has proven an excellent screen for regeneration treatments. Promoting retinal ganglion cell regeneration has the potential to restore vision loss associated with optic neuropathies such as glaucoma, and virally delivered gene therapy for eye disease is already in clinical practice (Bainbridge *et al*., 2008). One of the most potent interventions in pre-clinical models up to date is a conditional deletion of the PTEN in adult RGCs which resulted in improved regeneration and neuronal survival after optic nerve crush. This effect was attributed to upregulation of the mTOR signaling pathway (Park *et al*., 2008). In the current study, adult mice were treated by delivering an AAV vector into the vitreous 2 weeks before an optic nerve crush. Both phosphomimetic active and wild-type Protrudin led to a large number of axons regenerating for a long distance only 2 weeks after optic nerve crush (**Fig. 5**). Expression of active Protrudin allowed for 400-500 neuronal fibers to reach the 0.5 mm mark by 2 weeks after injury suggesting that this intervention is comparable to the most potent interventions reported to date.

In addition, Protrudin’s overexpression (both wild-type and active forms) was completely neuroprotective in a retinal explant model of RGC injury (**Fig. 6**). This effect could be due to increased signaling as a result of improved axonal transport which in turn activates retrograde survival signals. Further studies are needed in order to pinpoint the exact mechanism of Protrudin-driven neuroprotection in the eye.

Our study demonstrates robust axon regeneration in the adult CNS driven by overexpression of the adapter protein Protrudin. Overexpression of Protrudin, particularly in its phosphomimetic, active form greatly enhanced regeneration in cortical neurons *in vitro* and in the injured adult optic nerve. Importantly, both Protrudin and active Protrudin expression lead to an accumulation of ER and enhanced axonal transport in the distal axon and interfering with Protrudin’s ER localization or transport domains abrogates its regenerative effects, indicating a central role for these processes in mediating Protrudin-driven regeneration. We propose that Protrudin enables regeneration by acting as a scaffold to link the ER, recycling endosomes, kinesin-based transport and membrane phospholipids. Our findings establish the importance of these components in facilitating CNS axon regeneration, whilst suggesting Protrudin gene-therapy as a potential approach for repairing CNS axon damage.

## Acknowledgments

We would like to thank Prof. Joost Verhaagen for providing the viral constructs in which Protrudin was cloned, Dr Andrew Osborne for supervising during eye injections, animal perfusions and tissue collections when validating the efficacy of the Protrudin viruses, Mr Tolga Sadku for creating the Protrudin schematic diagram, the Light Microscopy Core of the NHLBI/NIH, and Mr. Raymond Fields at the National Institute of Neurological Diseases and Stroke Viral Core Facility for making the Protrudin viruses.

## Author contributions

Conceptualization, V.P., R.E., E.R. J.W.F.; Methodology, V.P., R.E., C.S.P., J.R.T, P.A.W., J.W.F.; Validation, V.P., C.S.P., A.S., J.R.T; Formal Analysis, V.P., C.S.P., A.S., J.R.T.; Investigation, V.P., C.S.P., A.S., J.R.T.; Data Curation, V.P., R.E., C.S.P., J.R.T., P.A.W.; Writing – Original Draft, V.P., R.E., J.W.F.; Writing – Review and Editing, all; Visualization, V.P., R.E., C.S.P., J.R.T., P.A.W; Supervision, E.R., P.A.W., H.M.G., R.E., J.W.F.; Funding Acquisition, V.P, P.A.W., H.M.G., R.E., J.W.F.

## Supplementary Information

**Fig. S1.**
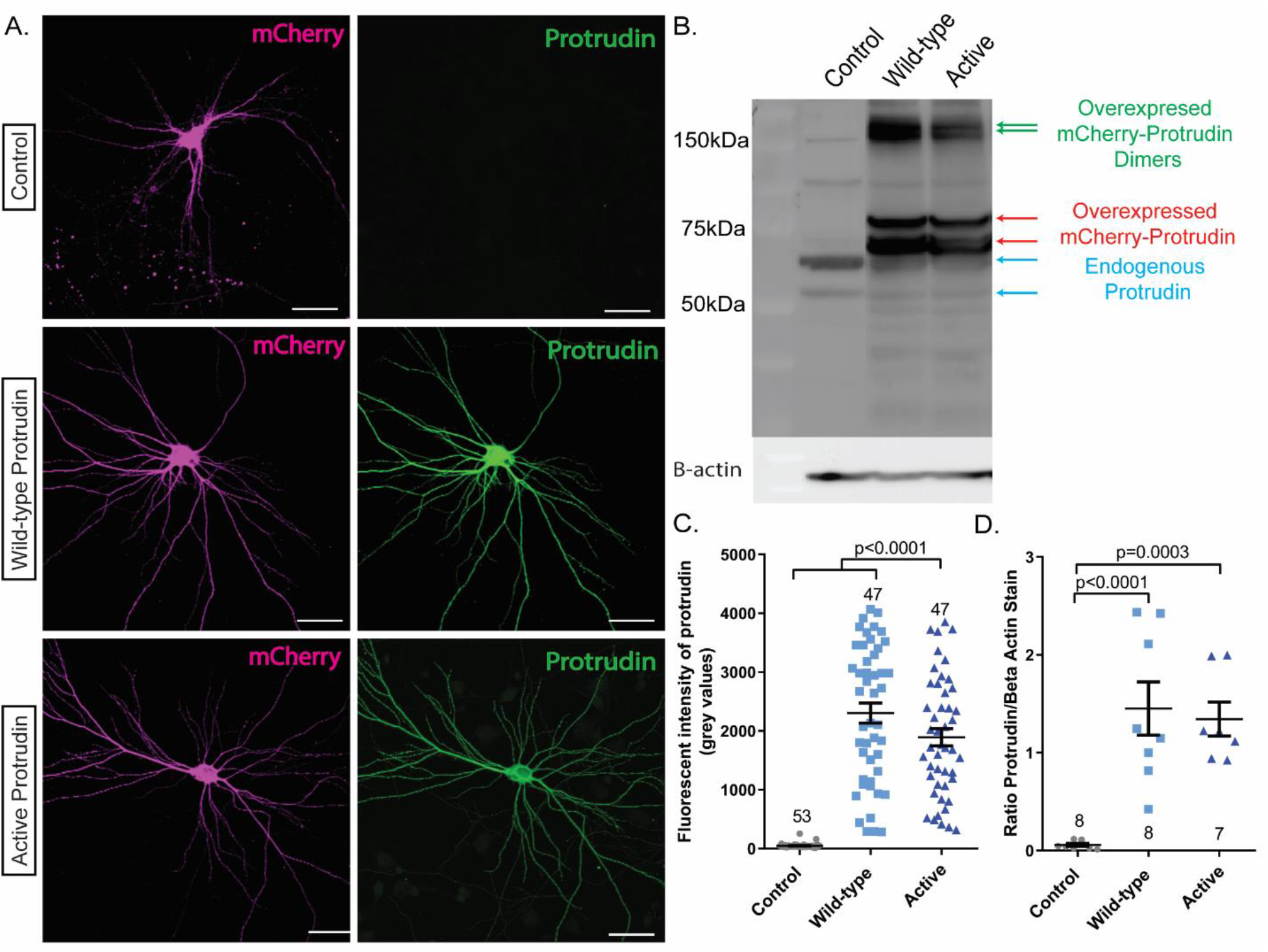
Overexpression of Protrudin results in higher amount of the Protrudin protein in rat cortical neurons at 14 DIV. (**A**) Images showing mCherry fluorescence (magenta) and Protrudin immunofluorescence (green) from neurons transfected with syn-mCh control (Control), syn-mCh-WT (wild-type Protrudin) or syn-mCh-active (active Protrudin) (magenta) and labelled with anti-Protrudin antibody. Scale bars are 20 μm. (**B**) Western blot showing Protrudin expression levels in PC12 cells overexpressing the three different constructs. The blot was stripped and probed for beta actin (lower panels). Blue arrows indicate endogenous Protrudin, red arrows indicate overexpressed Protrudin-mCherry and green arrows show overexpressed mCherry-tagged Protrudin dimers. (**C**) Scatter plot to show the average fluorescence intensity of Protrudin in transfected neurons (*n* = 3, *Kruskal-Wallis with Dunn’s* multiple comparison test, *p* < 0.0001). (**D**) Scatter plot to show the ratio of protrudin to beta actin staining of PC12 lysate extracted from cells overexpression either control, wild-type protrudin or active protrudin constructs (*n* = 7-8, *Kruskal-Wallis with Dunn’s* multiple comparison test, *p* < 0.0001). Error bars represent mean ± SEM.

**Fig. S2.**
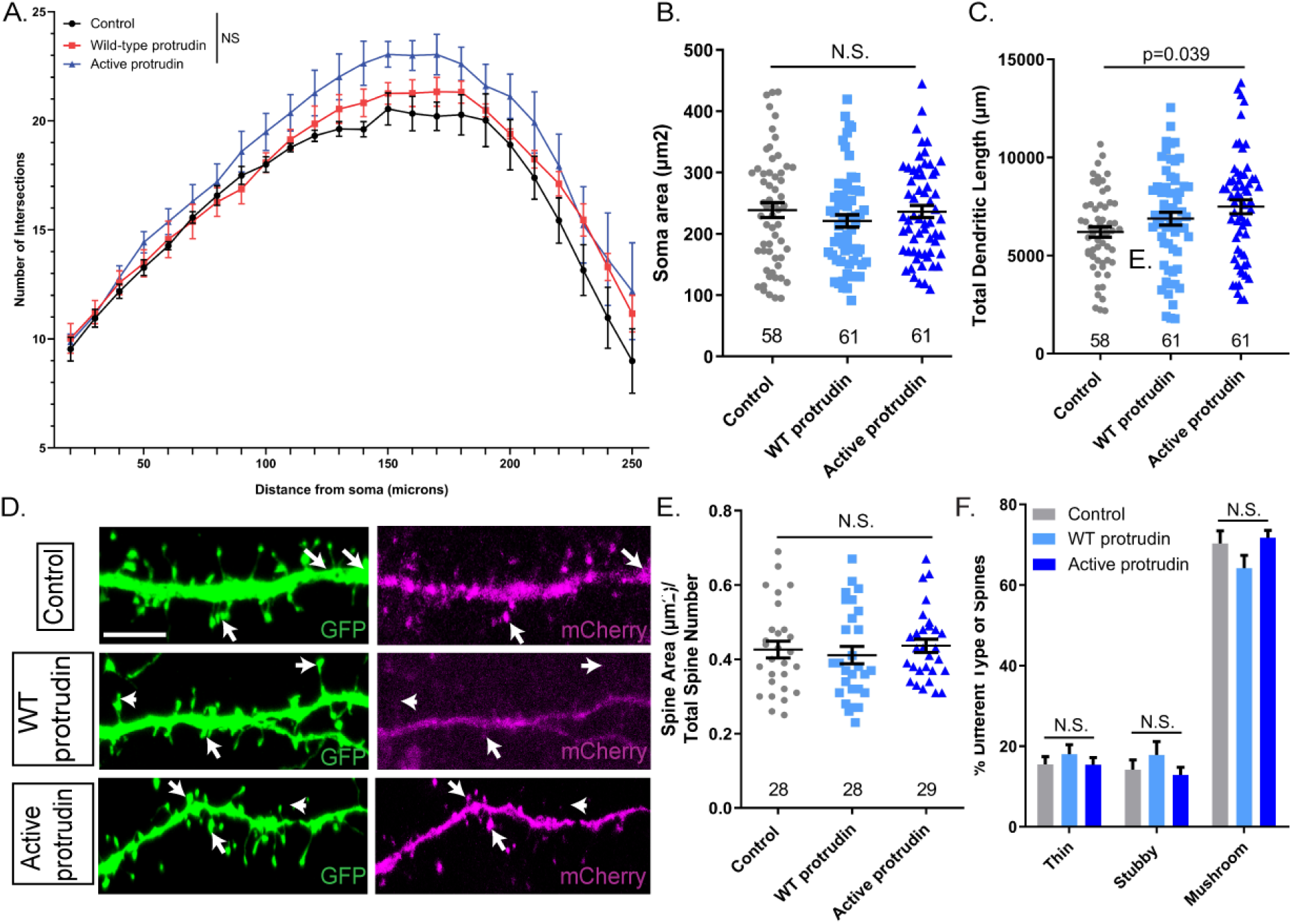
Overexpression of Protrudin does not result in morphological changes. (**A**) Dendritic tree complexity of neurons transfected with each construct - the number of intersections at each distance from the cell soma was plotted for each condition. There are no significant differences between the conditions (*n* = 4, repeated measures *2-way ANOVA*). (**B**) Average soma area in cells expressing the three different mCherry constructs (*n* = 5, *p* = 0.142, *Kruskal-Wallis with Dunn’s* multiple comparison test). (**C**) Dendritic tree total length across the different conditions. Cells overexpressing active Protrudin have a more complex dendritic structure than cells overexpressing control (*n* = 4, *p* = 0.067, *one-way ANOVA*). (**D**) Representative z-project images of 20 μm z-stack sections examined for dendritic spine number and morphology. White arrows point to individual spines. Scale bars are 5 μm. (**E**) The spine area (μm^2^) per the total number of spines for each condition. There were no significant differences between the four conditions (*n* = 2, *p* = 0.127, *Kruskal-Wallis with Dunn’s* multiple comparison test). (**F**) There are no changes in spine morphology between the different conditions.

**Fig. S3.**
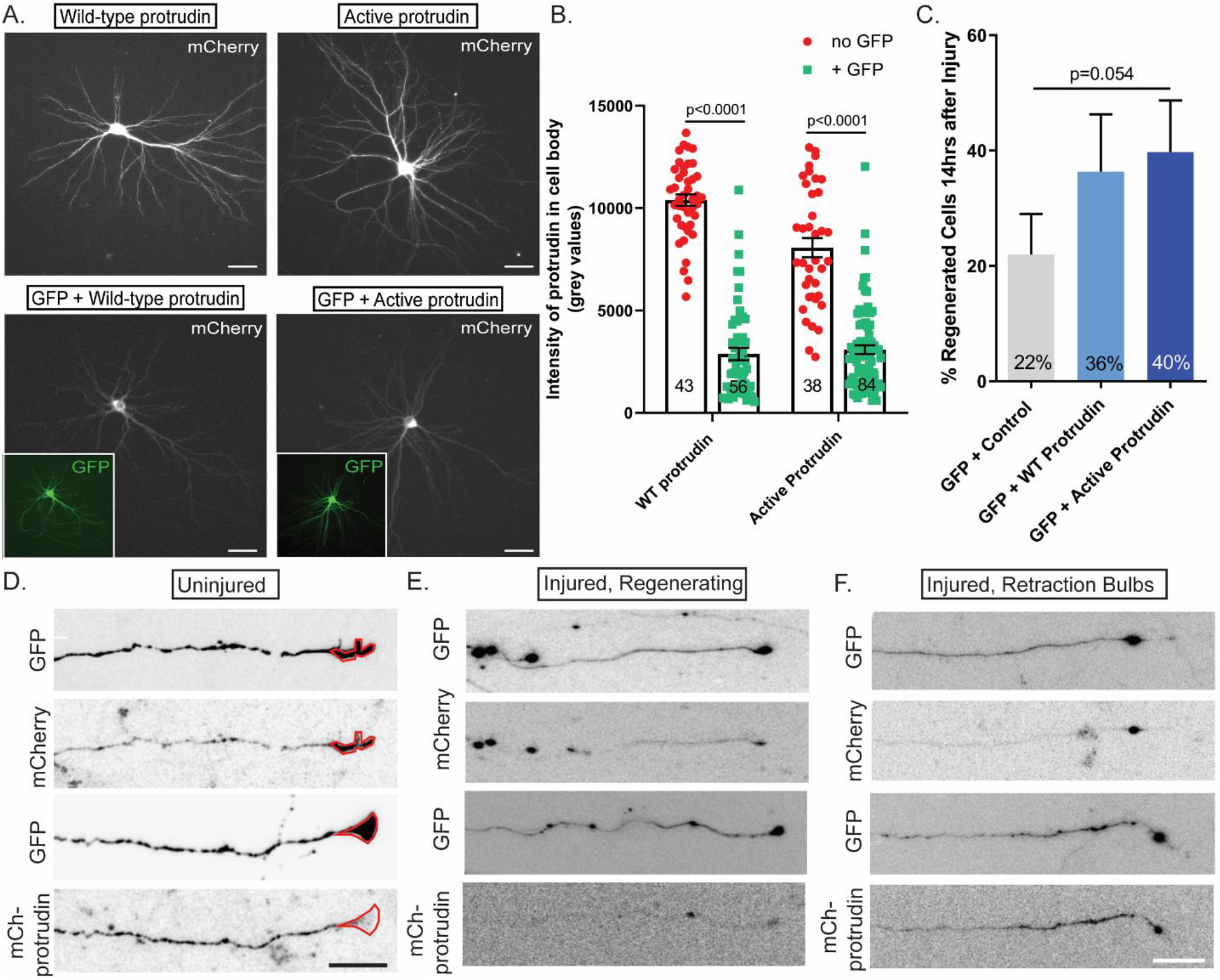
Protrudin promotes regeneration in a dose-dependent manner and localizes to the axon tip. (**A**) Immunofluorescence images of cells expressing wild-type or active Protrudin on their own or in combination with GFP control plasmid. Scale bars are 20 μm. (**B**) Quantification of mCherry-Protrudin fluorescence intensity in the soma. (**C**) Percentage of regenerating axons after laser axotomy in neurons co-transfected with Protrudin constructs and GFP. (**D**) Representative images showing GFP and mCherry-Protrudin at the growth cone of neurons co-expressing both constructs. Red lines indicate the area of the growth cone as defined by the GFP signal. Scale bars are 10 μm. (**E**) Images showing regenerating axons after injury. Scale bars are 20 μm. (**F**) Images showing retraction bulbs of non-regenerating processes after laser axotomy. Scale bars are 20 μm.

**Fig. S4.**
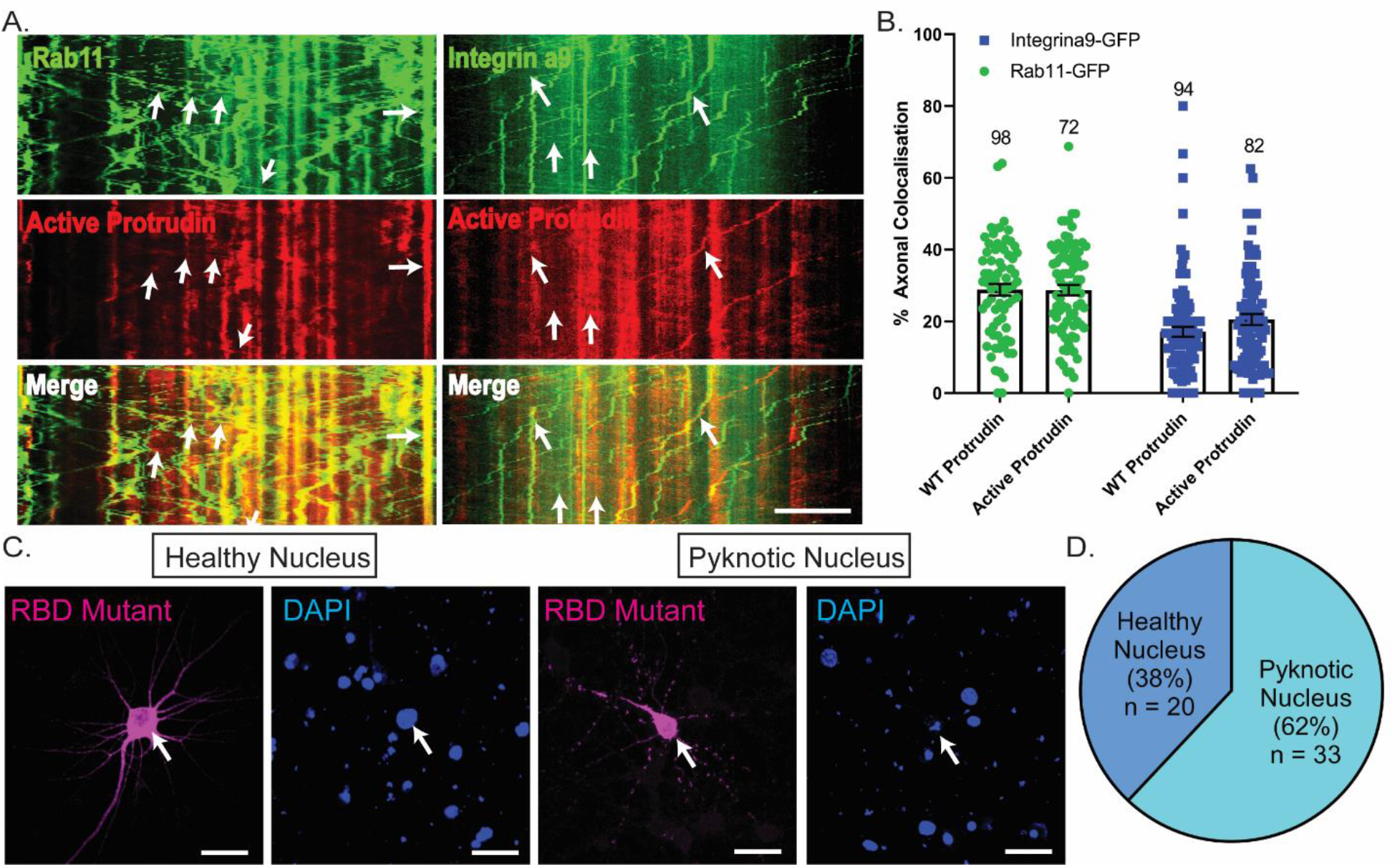
Rab11 and integrin vesicles co-localize with Protrudin and RBD domain deletion has consequences on cell survival. (**A**) Kymographs showing co-localization between Rab11 or integrin α9-GFP with Protrudin. (**B**) Wild-type and active Protrudin co-localize to similar extent with either Rab1 or integrin alpha 9. Error bars represent mean ± SEM. (**C**) Representative fluorescent images of RBD mutant of Protrudin in surviving and non-surviving (the most common phenotype) cells as shown by nuclear fragmentation (pyknosis). Scale bars are 20 μm. (**D**) Pie chart showing the percentage of cells with healthy and pyknotic nuclei in cortical neurons overexpressing RBD mutant protrudin (*n* = 3).

**Fig. S5.**
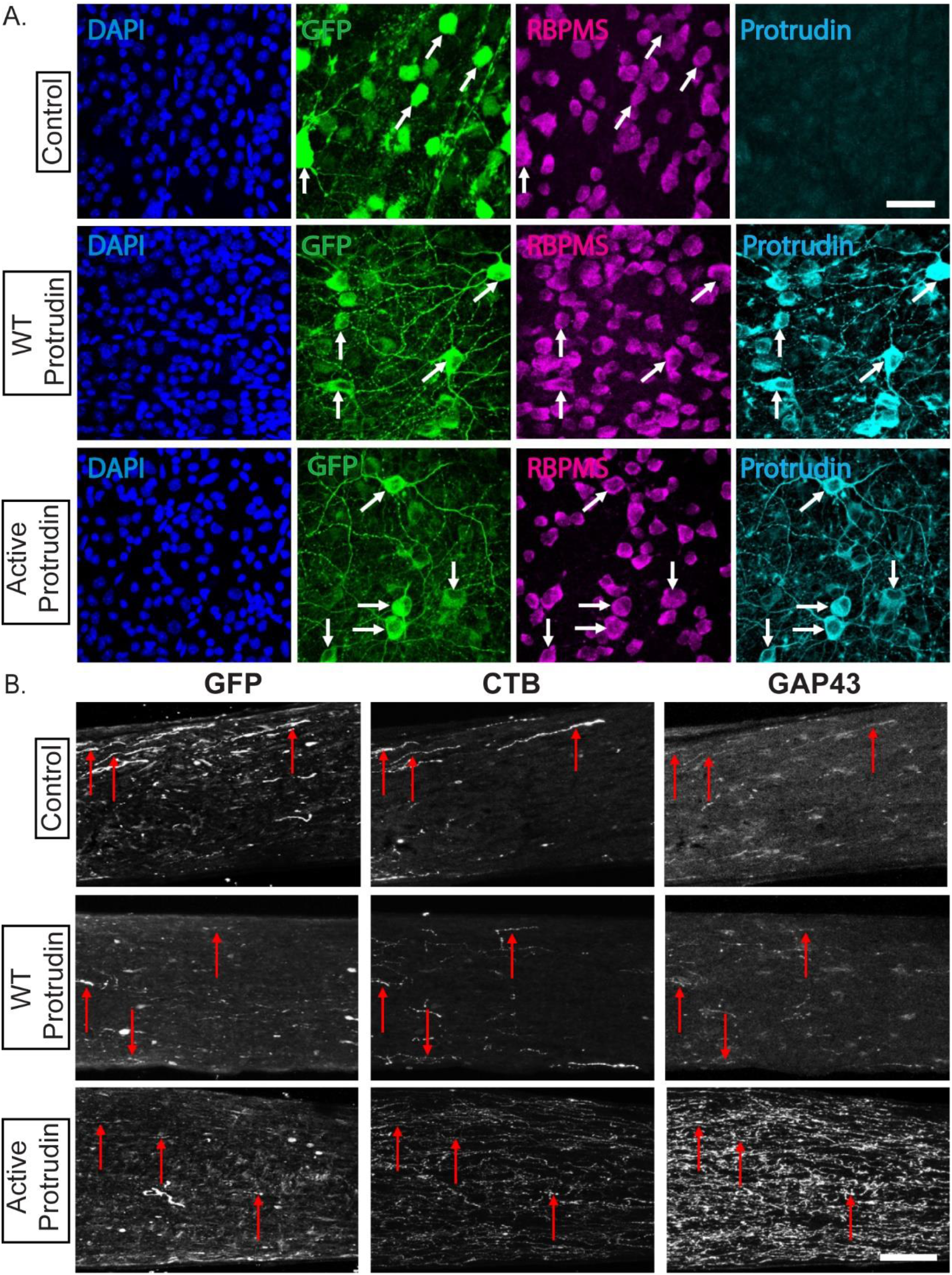
Protrudin viruses are expressed in RGCs throughout the retina and the optic nerve and regenerating axons are GAP43 and CTB positive. (**A**) Retinal sections showing co-localization between RBPMS (magenta) and GFP (green) and elevated Protrudin levels (cyan) as detected by immunohistochemistry. Scale bars are 20 µm. (**F**) Immunolabelled optic nerve sections for GAP-43, CTB and GFP. Red arrows show regenerating axons where GFP, CTB and GAP43 all co-localize. Scale bars are 10 µm

**Movie S1.** Example videos showing a non-regenerating axon expressing mCherry control plasmid and regenerating axons expressing either wild-type or active protrudin after laser axotomy over a 14-hour period of imaging.

### Contact for Reagent and Resource Sharing

Further information and requests for resources and reagents should be directed to and will be fulfilled by the Lead Contact Veselina Petrova.

### Experimental Models and Subject Details

#### Animals

All procedures were performed in accordance with protocols approved by the Institutional Animal Care and Use Committee (IACUC) at the National Institutes of Health, by the UK Home Office regulations for the care and use of laboratory animals under the UK Animals (Scientific Procedures) Act (1986) and in accordance with the Swedish Board of Agriculture guidelines and were approved by the Karolinska Institutet Animal Care Committee. Female C57Bl/6 mice aged 6-8 weeks (Charles River) were housed in a pathogen-free facility with free access to food and a standard 12 h light/dark cycle. Female pregnant Sprague Dawley rats were obtained from Charles River Laboratories, and embryos (both genders) at E18 stage of development were used for primary cultures of cortical neuron.

#### Primary neuronal cultures and transfection

Rat cortical neurons were dissected from E18 embryos and plated on imaging dishes or on acid-washed glass coverslips at the following densities: 1×10^5^cells/dish for immunocytochemistry, 2×10^5^ for axotomy or live-cell imaging and 8×10^4^ cells/coverslip. All surfaces were coated with poly-D-lysine, diluted in borate buffer to a final concentration 50 μg/mL. The cells were grown in serum-free MACS Neurobasal Media supplemented with 2% NeuroBrew21 and 1% GlutaMAX supplements at 37°C in 7% CO2 incubator. Cortical neurons were transfected using NeuroMag magnetofection system where 7μg of DNA plasmid is mixed with 100μL NB media and 8μL of magnetic beads. The reaction was kept for 30 minutes at 37°C before adding 900μL of pre-warmed NB media to a final volume of 1 mL. The original neuronal media was removed, and 1 mL of transfection mixture was added. Dishes were then incubated at 37°C for 30 minutes on a magnetic plate before the original media was returned on the plates. Plasmid reporter gene expression was observed 48 hours post-transfection.

#### Methods Details

##### DNA constructs

Human Protrudin constructs (in pmCherry-C1 and pEGFP-C1), CMV-integrin-alpha9-GFP and CMV-Rab11-GFP were used. The CMV promoter in all constructs was replaced by a human synapsin (Syn) promoter by Gibson assembly cloning. The viral vector plasmid backbones (AAV2-sCAG-GFP) were a kind donation by Prof. Joost Verhaagen, The Netherlands Institute for Neuroscience. Protrudin-GFP was cloned from pEGFP-C1 plasmid into viral vector plasmids using Gibson cloning. Site-direct mutagenesis was performed in order to create the Protrudin active phosphomimetic form (QuikChange II Site-Directed Mutagenesis Kit, Agilent Technologies). All constructs were verified by DNA sequencing.

##### Immunostaining

Cortical neurons were fixed in 4% PFA for 15 minutes and then thoroughly washed and kept in PBS at 4°C. Cells were permeabilized in 3% BSA in PBS and 0.1% Triton for 5 minutes and then blocking solution was added (3% BSA in PBS) for 1 hour at room temperature. Primary antibodies were added at the correct concentration and kept for 1.5 hours at room temperature. Antibodies were then washed 3 times in PBS for 5 minutes. Secondary antibodies were applied at the correct concentration for 1 hour at room temperature in a dark chamber. The cells were then washed three times in 1xPBS and mounted using coverslips and Diamond anti-fade mounting agent with DAPI (Molecular Probes) or FluorSave mounting reagent (Calbiochem). Mice were anesthetized using 1-2% isoflurane and transcardially perfused with PBS followed by 4% paraformaldehyde (PFA). Optic nerves were dissected and immersed in 4% PFA. The tissue was post-fixed overnight, then immersed in 30% sucrose for 24 h for cryoprotection. Tissue was embedded in Tissue-Tek OCT and snap-frozen for cryosectioning. 14 μm-thick longitudinal sections of the optic nerve were obtained on charged Superfrost microscope slides using a Leica CM3050 cryostat. Slides were dried and stored at −20°C.

##### Live-cell imaging

Live-cell imaging was performed using spinning disk confocal microscopy, using an Olympus IX70 microscope with a Hamamatsu EM-CCD Image-EM camera and a PerkinElmer Ultra-VIEW scanner. Videos were taken using Meta-Morph software. Rab11 and integrin vesicle trafficking along the axon was imaged at the proximal (up to 100μm) and distal part (beyond 600μm) of axons of neurons transfected with Protrudin (as described before in Eva *et al*., 2010, 2017). One image per second was obtained for 3 minutes. Kymographs were obtained by measuring an individual axon segment. Anterograde, retrograde, bidirectional and static modes of transport were measured. The percentage of co-localization between integrin or rab11 and Protrudin was calculated as the number of vesicles containing either was divided by the total number of vesicles. All analysis was performed using Meta-Morph software.

##### Western Blotting

PC12 cells were transfected using lipofectamine. 48 hours later cells were lysed using the cOmplete Lysis Kit (Roche). Cells were washed with ice-cold PBS. 500 μL of pre-cooled lysis buffer was added to each well of a 6-well plate and the lysate was scraped using a cell scraper and transferred to a sterile 1.5 mL Eppendorf. The lysate was incubated on ice for 30 minutes with occasional mixing. The samples were then centrifuged at 10 000 g at 4°C for 10 minutes. The supernatant was transferred to 1.5 mL Eppendorf and the pellet was discarded. The total protein concentration was measure by BCA assay using Pierce BCA Assay Kit Protocol (ThermoFischer Scientific). The 96-well plate containing the sample lysates and BCA reagents was analyzed using Gen5.1 software and concentrations were derived from a standard curve for albumin control. 15 μg of PC12 cell lysate was then treated with LDS Sample Buffer NuPAGE 4x (1:4, ThermoFisher Scientific) and Sample Reducing Agent (1:10, ThermoFisher Scientific) and were analyzed by Western blot. Samples were run on a 4-12% gel at 120 V at room temperature in 40 mL Running Buffer NuPAGE (ThermoFisher Scientific) diluted in H20 to 800 mL. The gel was then transferred onto a nitrocellulose membrane (Invitrolon PDVF/Filter Paper Sandwich, ThermoFischer Scientific) for 1.5 hours at 40V at 4°C in 50 mL Transfer Buffer NuPAGE (ThermoFisher Scientific) in 100 mL methanol, topped up with ddH20 to 1 L. The membrane was then blocked either in 5% milk or 3% BSA depending on the antibody for 1 hour and incubated overnight with the primary antibody diluted the blocking solution to the right concentration at 4°C. The membrane was then rinsed three times in Tris-buffered saline with Tween 20 (TBST buffer) for 10 minutes each. The TBST buffer was removed. Secondary peroxidase-conjugated antibodies were diluted to the right concentration in blocking solution and were then added for 1 hour at room temperature. SuperSignal West Dura Extended Duration Substrate kit (ThermoFischer Scientific) and Alliance software were then used for detection.

##### Laser axotomy

10DIV neurons were transfected with various constructs using magnetofection as described above. Between 13-17 DIV, their regeneration ability was examined using the laser axotomy model described in detail in previous reports. Axotomy was performed by an UV Laser (355 nm, DPSL-355/14, Rapp OptoElectronic, Germany) connected to a Leica DMI6000B (Leica Systems, Germany), and all images were taken with an EMCCD camera (C9100-02, Hamamatsu). Axons were cut at least 600 μm away from the cell body and regeneration was observed for 14 hours post injury at 30-minute intervals. If more than 50% neuronal cell death occurred in the axotomised cells, the experiment was excluded from the final analysis.

##### Animal Studies

Intravitreal injections of viruses were administered 14 days prior to optic nerve crush or whole retinal explant. 5 μL of the injecting solution for mice was drawn into a sterile 5 μL Hamilton syringe equipped with a 33-gauge removable needle. The solution was injected into the vitreous humor through the superior nasal sclera, with the needle positioned at a 45° angle to avoid the lens, external ocular muscle, and blood vessels. Before removing the needle, a sterile 33-gauge needle was used to puncture the cornea and drain the anterior chamber, thereby reducing intraocular pressure and preventing reflux. Different needles were used for each virus to prevent contamination, and syringes were rinsed between injections with ethanol followed by sterile PBS.

##### Optic Nerve Crush

Optic nerve crush was performed as described in Pearson *et al*., 2018. Micro-scissors were used to make an incision in the conjunctiva and expose the optic nerve. Curved forceps were then inserted below the external ocular muscle, avoiding the ophthalmic artery and retrobulbar sinus, and positioned around the exposed nerve. The nerve was crushed for 10 s approximately 1 mm behind the eye. Following the crush, eyes were observed fundoscopically for signs of ischemia, and mice were monitored for signs of intraorbital bleeding. Mice were given a subcutaneous injection of 1 mg/kg buprenorphrine as an analgesic and topical application of ophthalmic ointment to prevent corneal drying. Intravitreal injections of CTB (1.0 μg/μL, Sigma) were administered 2 d prior to perfusion. 2 μL of the solution injected as described above.

##### Whole retina explant culture

Mice were euthanized by cervical dislocation, eyes enucleated, and placed immediately into ice-cold HBSS. Retinas were dissected from the eyes in HBSS on ice, flat-mounted with the ganglion cell layer up on a cell culture insert (Millipore), and submerged in tissue culture media containing Neurobasal –A, 1% penicillin-streptomycin (10000 U/ml), 1% glutamine (100X), 1% N-2 (100X) and 1% B-27 (50X) (all ThermoFisher Scientific). Retinas were incubated in 6-well plates at 37 °C and 4% CO_2_ for 3 days and were fed by replacing 50% of the media on day 2. Retina were fixed in 3.7% PFA and stained with antibodies against RBMPS and GFP followed by counterstaining with DAPI. For untreated “DEV 0” controls, retinas were dissected and placed straight into 3.7% PFA.

##### Statistics

Statistical analysis was performed using GraphPad Prism 8.0 (GraphPad Software, La Jolla, CA). Each data set was individually tested for normal distribution using the D’Agostino-Pearson normality test. When data was normally distributed one-way ANOVA with multiple comparisons was used to test statistical significance between the experimental groups with Tukey’s post hoc test. Several data sets were shown to be non-normally distributed. Therefore, a non-parametric Kruskal-Wallis test with multiple comparisons was used to test for significant differences across experimental groups. Dunn’s multiple comparison post-hoc test was also performed. All statistics were carried out at 95% confidence intervals, therefore a significant threshold of p<0.05 was used in all analyses. For Sholl analysis, repeated measures two-way ANOVA was performed using SPSS (IBM Statistics). When comparing percentages (e.g. of regenerating cells), Fisher’s exact test was performed between each two groups compared. p-values were then analyzed with the “Analyze a stack of p values” function in GraphPad Prism with a Bonferroni-Dunn pairwise comparison to test for statistical significance between groups.

### KEY RESOURCES TABLE

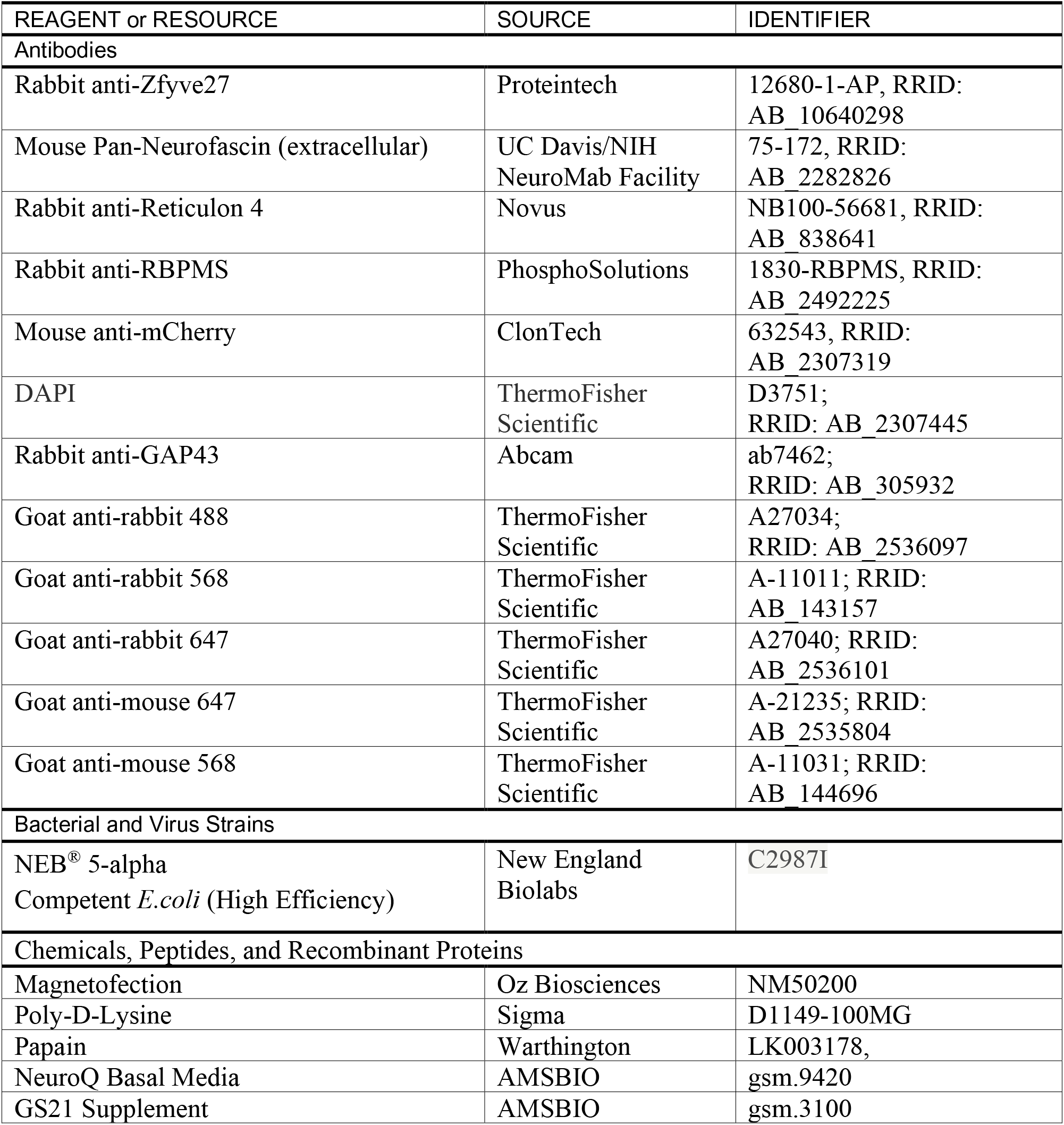
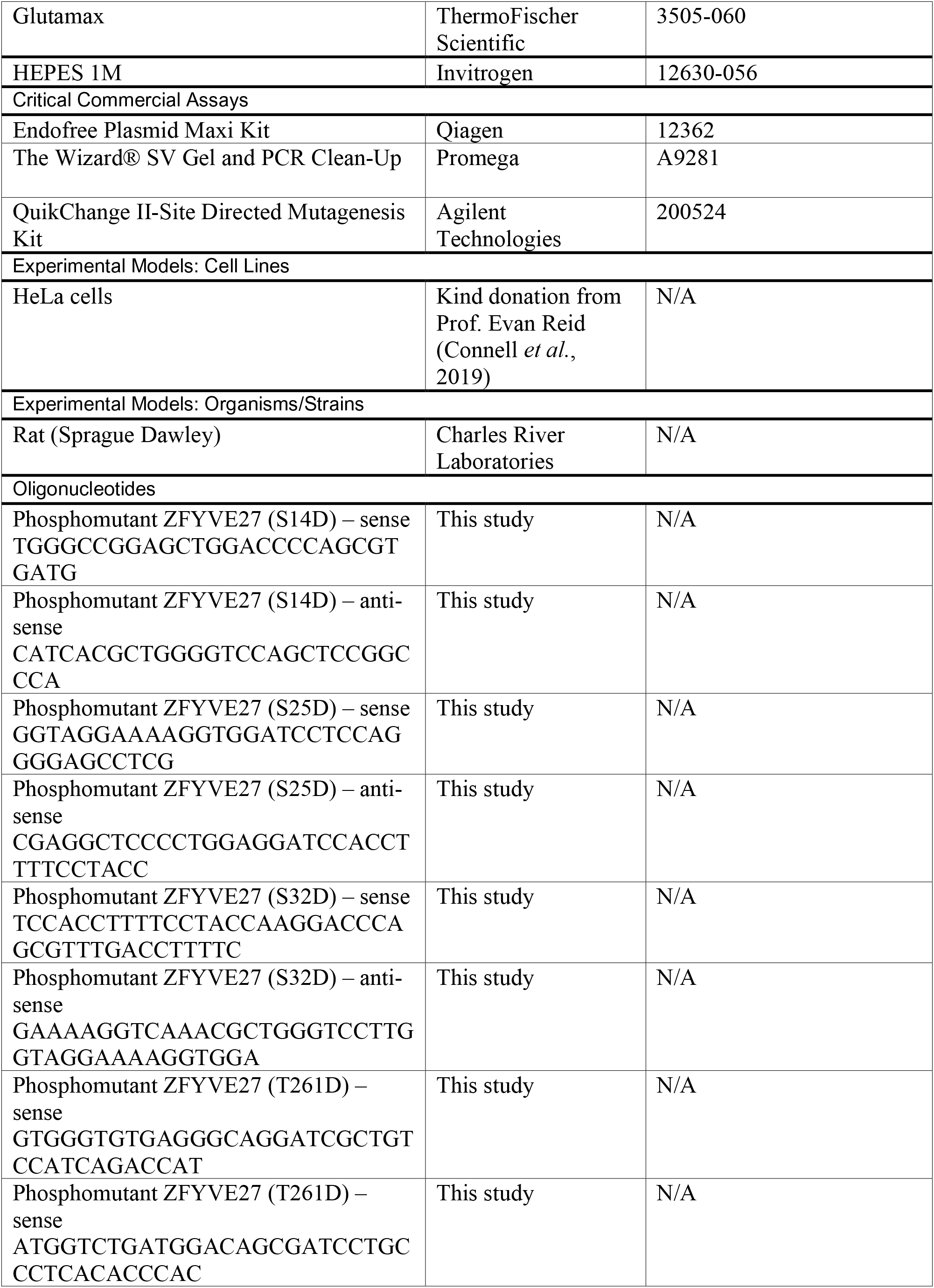
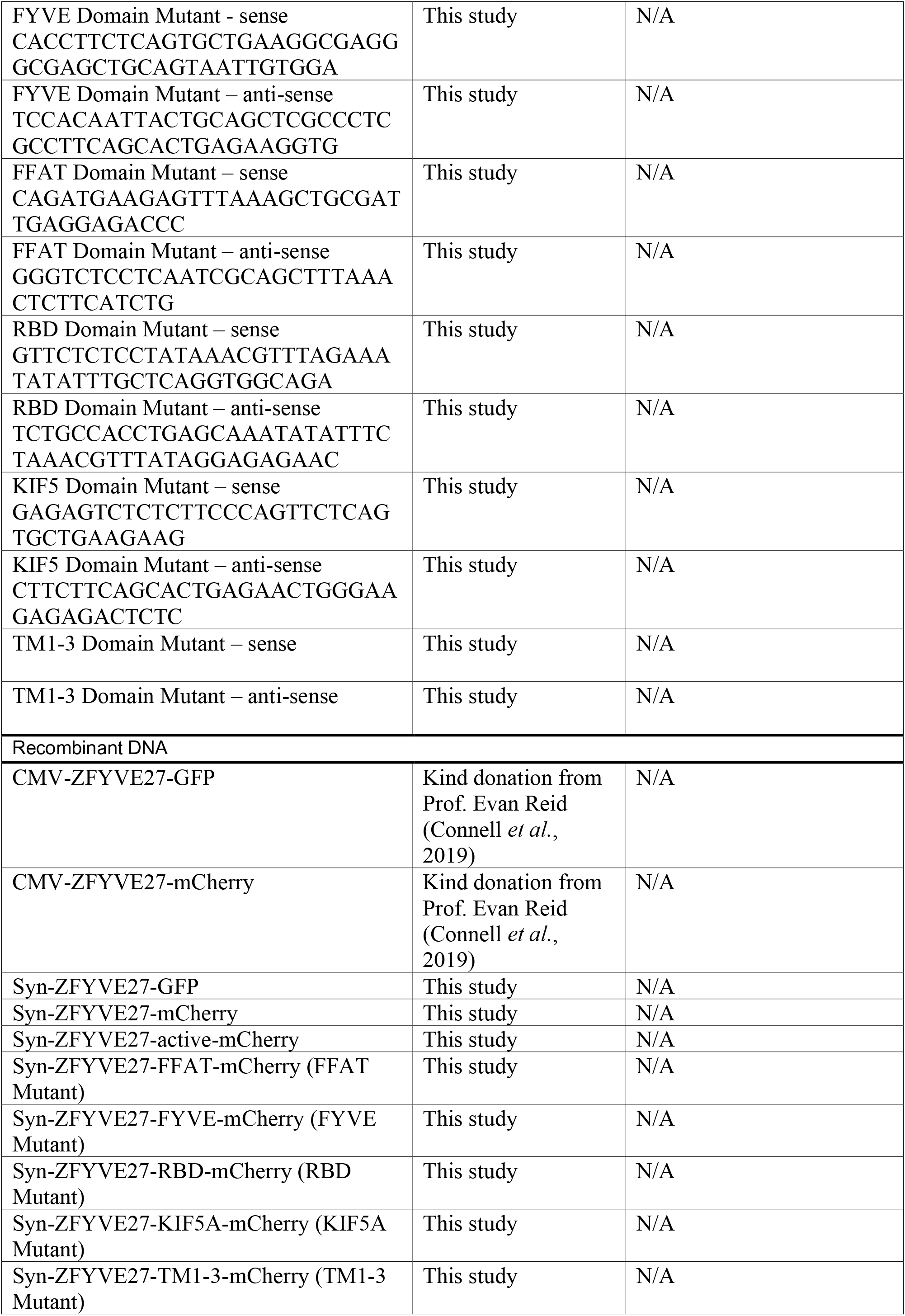
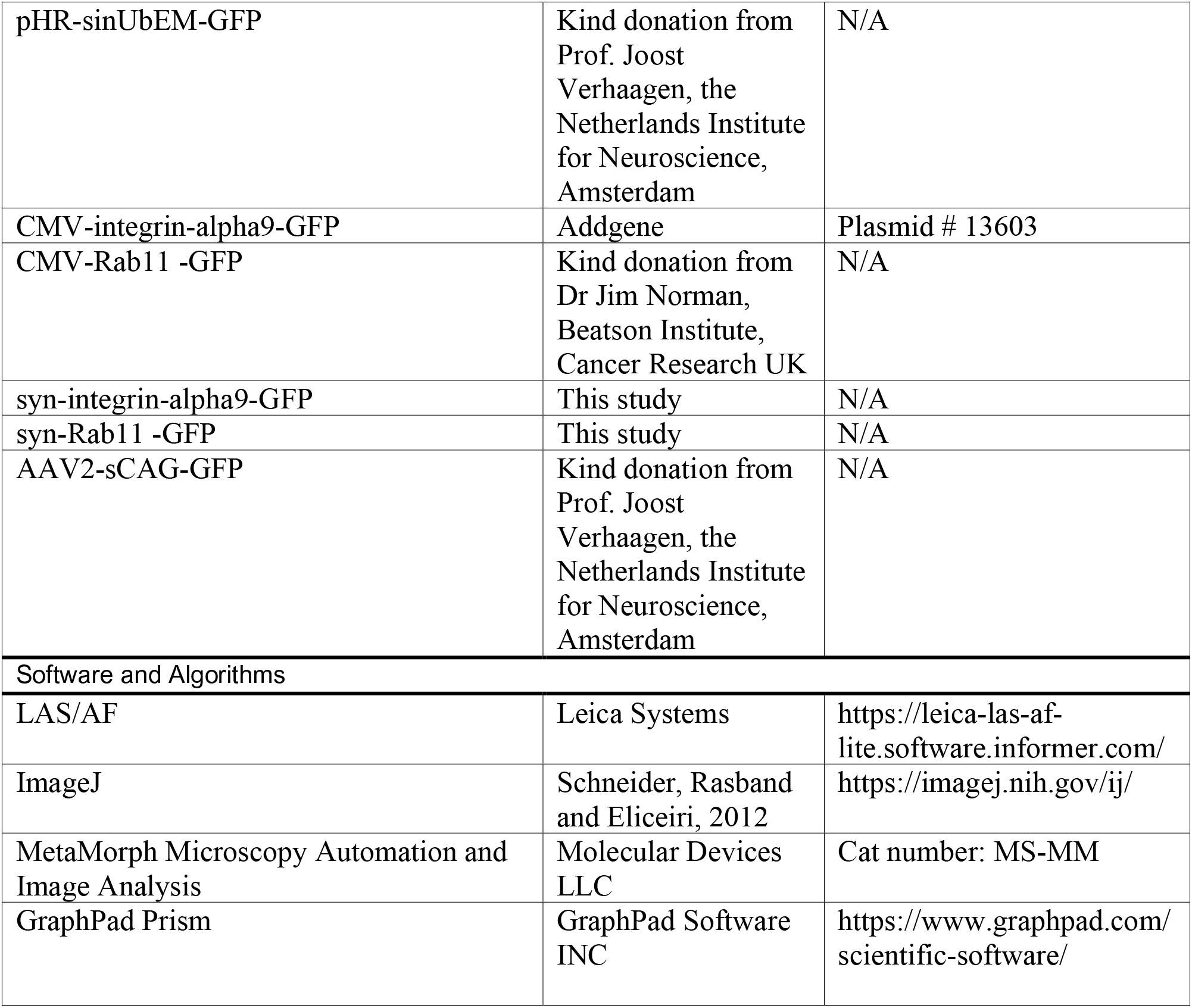

